# Cytoplasmic protein-free mRNA induces stress granules by two G3BP1/2-dependent mechanisms

**DOI:** 10.1101/2024.02.07.578830

**Authors:** Sean J. Ihn, Laura Farlam-Williams, Alexander F. Palazzo, Hyun O. Lee

## Abstract

Stress granules are cytoplasmic membraneless organelles that sequester proteins and non-translating mRNAs in response to various stressors. To assess the contributions of mRNA and RNA-binding proteins to stress granule formation, we use microinjection to deliver protein-free mRNA into the cytoplasm in a controlled manner. We demonstrate that mRNAs trigger stress granule formation through two mechanisms that are enhanced by the presence of G3BP1 and G3BP2. Low concentrations of *in vitro* transcribed mRNA activated protein kinase R (PKR), leading to phosphorylation and inhibition of the eukaryotic translation initiation factor eIF2α and stress granule formation. This was inhibited by replacing uridine with pseudouridine in the mRNA or by treating it with RNase III, which cleaves double-stranded RNA. High concentrations of mRNA triggered stress granule formation by a mechanism that was independent of PKR and enhanced by G3BP1/2, highlighting the importance of both protein-free mRNA and RNA-binding proteins in stress granule formation.

**Graphical Abstract/Model:** 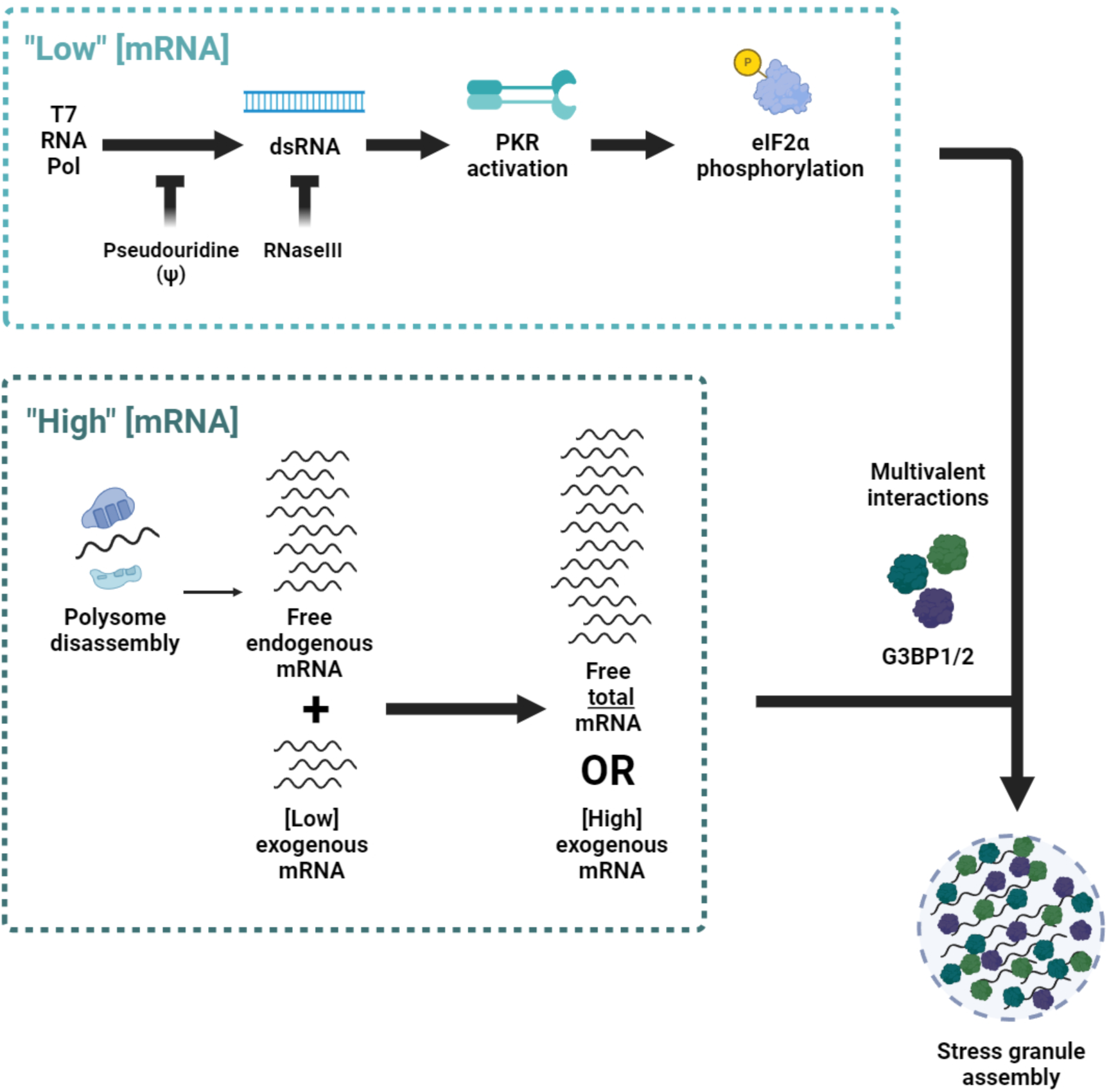

**Summary:** Microinjected mRNA induces stress granules in mammalian cells by two G3BP1/2-dependent mechanisms: one requires the stress-sensing protein kinase PKR to phosphorylate the translation initiation factor eIF2α, and the other is independent of PKR and phospho-eIF2α and acts when the cytoplasmic concentration of ribosome-free mRNA is increased acutely.

## Introduction

Stress granules are cytoplasmic biomolecular condensates, non-membrane-bound organelles that concentrate certain molecules and exclude others (Banani et al. 2017), that form in eukaryotic cells in response to a wide range of stressors, including oxidative stress, heat shock, osmotic stress and viral infection (Kedersha and Anderson, 2007; Hofmann et al., 2021; Buchan and Parker, 2009; Spannl et al., 2019). They concentrate various RNA-binding proteins, non-translating mRNAs, and nuclear transport factors and are proposed to regulate translation and coordinate nucleocytoplasmic transport during cell stress (Decker and Parker, 2012; Kedersha et al., 2013; Zhang et al., 2018). Stress granules are implicated in various disease states, including increased cancer cell survival (Fonteneau et al., 2022; Yang et al., 2022; Shi et al., 2019) and formation of toxic protein aggregates in amyotrophic lateral sclerosis and frontotemporal dementia, Huntington’s disease, Alzheimer’s disease and other neurodegenerative proteinopathies (Wolozin and Ivanov, 2019; Sanchez et al., 2021; Gutiérrez-Garcia et al., 2023; Baradaran-Heravi et al., 2020; Ash et al., 2014; Yu et al., 2022; Chew et al., 2019), which underscores the importance of understanding their regulation. Accordingly, the protein and RNA components of stress granules have been extensively studied, but their relative contribution to stress granule regulation is not well understood.

Stress granule formation is closely associated with global translation arrest and polysome disassembly, which liberates ribosome-bound mRNAs into the cytosol. Stressors that induce phosphorylation of translation initiation factor eIF2α block translation initiation and thereby trigger dissociation of mRNA from ribosomes, which coincides with stress granule formation (Kedersha et al., 1999). Conversely, freezing ribosomes on mRNAs by using inhibitors of translation elongation prevents stress granule assembly in mammalian cells under oxidative stress (Kedersha et al., 2000; Panas et al., 2016; Bounedjah et al., 2014). Accordingly, it has been proposed that excess ribosome-free mRNAs in the cytosol drive stress granules formation through multivalent, weak RNA–RNA interactions (Van Treeck and Parker, 2018) that lead to their phase separation, or demixing from the surrounding millieu to form biomolecular condensates (Hyman et al., 2014). Supporting this model, protein-free mRNAs purified from yeast readily phase separate in the presence of polycations *in vitro* (Van Treeck et al., 2018). Furthermore, treating cells with endonucleases specific for ssRNA blocks stress granule assembly under oxidative stress (Sanders et al., 2020) and leads to the dissolution of pre-formed stress granules (Guillén-Boixet et al., 2020), highlighting the importance of RNAs in stress granule assembly and maintenance. Whether excess ribosome-free mRNA in the cytosol is sufficient to drive stress granule formation, however, is not known.

Despite the importance of mRNAs in stress granule formation, it is their protein components that have been most thoroughly studied, some of which play crucial roles in stress granule formation. Depletion or deletion of the RNA-binding proteins G3BP1and G3BP2, or the RNA-binding protein TIA-1 blocks or significantly diminishes stress granule formation under most stress conditions tested (Protter and Parker, 2016; Gilks et al., 2004; Waris et al., 2014; Matsuki et al., 2013). Overexpression or induced oligomerization of G3BP1 triggers stress granule formation in the absence of stress (Reineke et al., 2012; Zhang et al., 2019), suggesting that G3BP1 functions as a scaffold for stress granule assembly. Furthermore, stress granule growth is facilitated by a network of G3BP1/2-interacting proteins (Youn et al., 2018; Cirillo et al., 2020; Kedersha et al., 2016) that are found in spatial proximity to each other even in the absence of stress (Youn et al., 2018). This raises the possibility that pre-existing interactions between stress granule proteins may nucleate the formation of stress granules. To what extent mRNAs and stress granule scaffolds like G3BP1/2 contribute to stress granule formation is unknown.

To assess whether an excess of ribosome-free mRNA in the cytoplasm can drive stress granule formation, we used microinjection as a means to introduce protein-free mRNA into the cytosol in a controlled manner. We report here that the microinjected mRNAs trigger formation of stress granules by protein kinase R (PKR)-dependent and -independent mechanisms. The PKR-dependent mechanism was activated even at low concentrations of microinjected mRNA and mediated eIF2α phosphorylation that could be blocked by the presence of a ubiquitous modified nucleotide, pseudouridine (5-ribosyluracil), or by treating the mRNA with RNase III, which selectively degrades double-stranded RNA (dsRNA). When PKR activity and eIF2α phosphorylation were blocked, microinjected mRNAs still triggered stress granule formation but only at high concentrations. The presence of scaffolding proteins G3BP1 and G3BP2 greatly enhanced stress granule formation by both the PKR-dependent and -independent mechanisms. Together, our findings demonstrate that cytosolic mRNAs drive stress granule assembly by two distinct mechanisms that are enhanced by scaffolding proteins G3BP1 and/or G3BP2 and highlight the importance of RNA binding protein-driven interactions in stress granule assembly.

## Results

### Microinjected mRNA triggers formation of stress granules and accumulates in them

To examine the role of protein-free mRNAs in stress granule assembly, we introduced mRNAs into human osteosarcoma (U2OS) cells by microinjection and examined stress granule formation at various timepoints. We used two model mRNAs: *fushi tarazu (ftz)* mRNA, which we previously found induces stress granules when microinjected into African green monkey kidney (COS-7) cells (Mahadevan et al., 2013), and *β-globin* mRNA. Both of these model mRNAs have been used extensively to study mRNA metabolism (Luo and Reed, 1999; Valencia et al., 2008). We transcribed these RNAs *in vitro* by using T7 RNA polymerase. To prevent their degradation, the mRNAs were capped and polyadenylated (we estimate the poly(A)-tail to be ∼100 nucleotides long; **Fig. 1 A**). The mRNA was co-injected with FITC-conjugated 70 kDa dextran, which does not cross the nuclear pore and thus marks the microinjected compartment. Each cell was microinjected with approximately 30–100 fl of mRNA at a concentration of 300 ng/μl, which we estimate corresponds to approximately 20,000–70,000 molecules per injection and 4–15% of the total amount of mRNA in the cell (around 500,000 mRNAs) (Carter et al., 2005). This had no significant effect on the total amount of mRNA in the cells, as determined by whole cell poly(A) staining following *ftz* mRNA microinjection (**Fig. 1 B, S1 A**).

**Figure 1.**
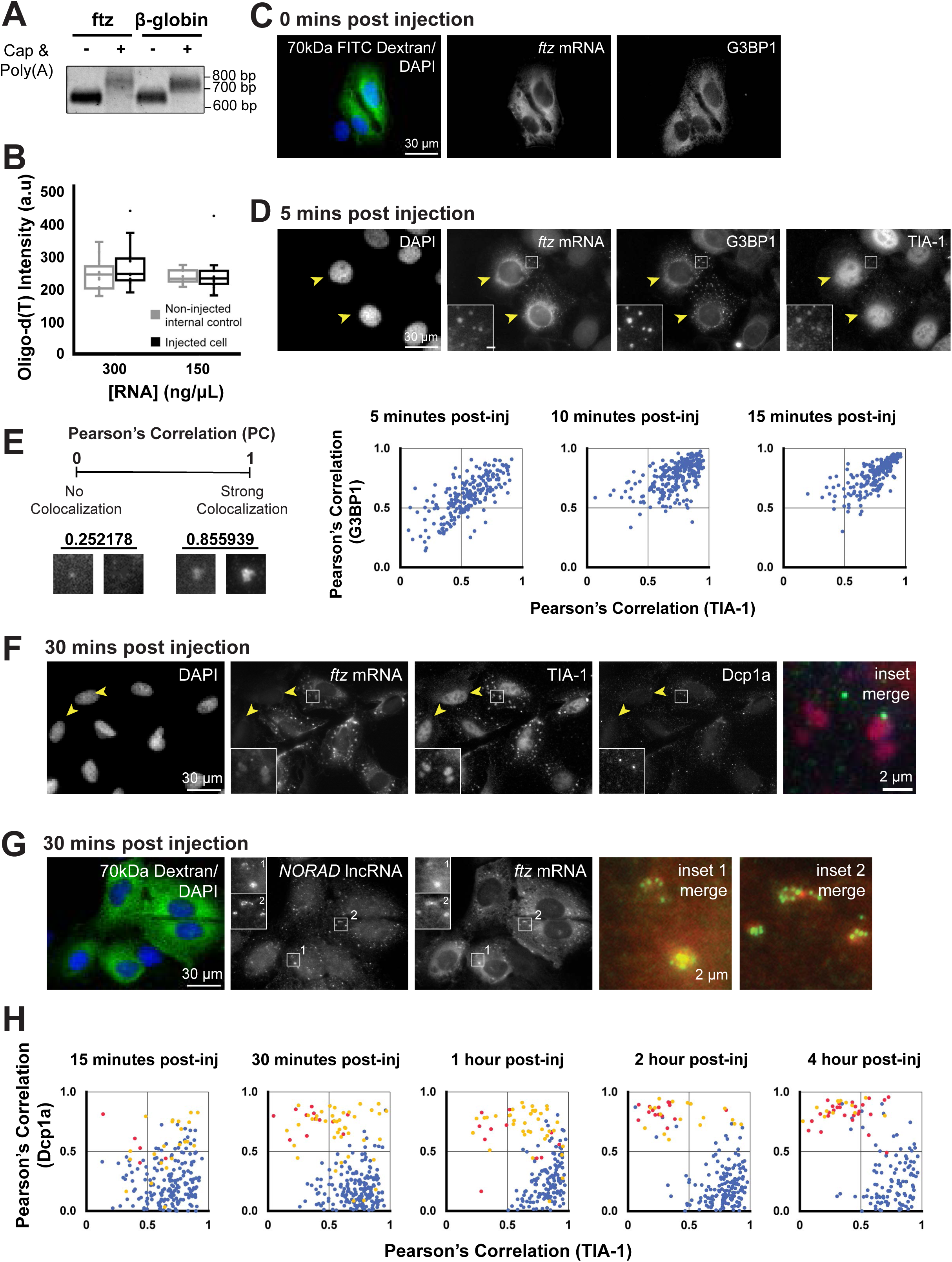
Microinjected mRNA triggers the formation of SGs and accumulates into them. (A) *ftz* and *β-globin* mRNAs were *in vitro* transcribed, capped and polyadenylated, then purified. Samples of mRNAs before (“-”) and after (“+”) capping and polyadenylation were collected and resolved on an a 1% agarose gel. The migration of standards are indicated on the right. (B) U2OS cells were microinjected with *ftz* mRNA (300 or 150ng/μL). 30 minutes after microinjection, the cells were fixed and stained against poly(A)-RNA by using oligo-dT probes. The whole cell oligo-dT staining intensities were quantified in arbitrary units (a.u.) between injected and non-injected cells. Each box plot displays the average, quartiles and standard deviations. (**C-D**) U2OS cells were microinjected with *ftz* mRNA (300ng/μL) and 70kD dextran conjugated to FITC to mark the microinjected cells and the compartment (cytoplasm). Immediately after microinjection (C) or after 5 minutes (D), the cells were fixed and stained against *ftz* mRNA using FISH probes, the SG-marker G3BP1 by immunofluorescence, and DNA by DAPI. Zoomed in images of boxed areas are shown in insets in (D). Scale bar=30µm. (E) U2OS cells were microinjected with *ftz* mRNA (300ng/μL) and 70kD dextran conjugated to FITC, incubated for the indicated times, fixed and stained for *ftz* mRNA, G3BP1 and TIA1. The cells were imaged and individual mRNA foci were evaluated for the colocalization of *ftz* mRNA with G3BP1 and TIA1 by Pearson Correlation Analysis. Examples of weak colocalization and strong colocalization are shown on the left. (F) U2OS cells were microinjected with *ftz* mRNA (300ng/μL). After 30 minutes the cells were fixed and stained with FISH probes against *ftz* mRNAs, the SG-marker TIA1, the PB-marker Dcp1a, and DAPI to stain DNA. All panels represent a single field of view imaged for the indicated components. Zoomed in images of inset (a) are displayed within boxes. The merged image show TIA-1 in red and Dcp1a in green. Scale bars: whole cell=30µm, insets=2µm. (G) U2OS cells were microinjected with *ftz* mRNA (300ng/μL) and 70kD dextran conjugated to FITC. After 30 minutes the cells were fixed and stained against *ftz* mRNA using FISH probes and endogenous *NORAD* lncRNA using single-molecule FISH probes, and DNA by DAPI. Scale bars: whole cell=30µm, insets=2µm. Zoomed in images are shown in boxed insets. The merge image shows *NORAD* lncRNA mRNA in green and *ftz* mRNA in red. (H) U2OS cells were microinjected with *ftz* mRNA (300ng/μL) and 70kD dextran conjugated to FITC. After 30 minutes the cells were immunostained for the SG-marker G3BP1 and the PB-marker Dcp1a. The cells were imaged and individual mRNA foci were evaluated for the colocalization of *ftz* mRNA with G3BP1 and TIA1 by Pearson Correlation Analysis. Foci that were small ad bright (see examples in Figure 1A-B) were labeled as PBs that were either docked to the surface of a SG (yellow), or a free floating (“undocked”; red). All other mRNA foci were labeled in blue. Note that most mRNA foci were positive for the SG marker and few resembled PBs that were docked to SGs at early timepoints. At later time points, the number of PBs increased and fewer of them were docked to SGs or contained G3BP1.

The cells were fixed at various times after microinjection and stained for *ftz* mRNA by using fluorescent *in situ* hybridization (FISH) and for the stress granule proteins G3BP1 and TIA-1 by immunofluorescence. Immediately after microinjection (0 minutes after injection), *ftz* mRNA, G3BP1 and TIA1 were located diffusely throughout the cytoplasm and nucleoplasm (**Fig. 1 C, S1 B**), indicating that there were no stress granules. Within 5 minutes, the *ftz* mRNA formed punctate structures in the cytoplasm that co-localized with G3BP1 and TIA1 (**Fig. 1 D**; microinjected cells are indicated by yellow arrowheads), consistent with the conclusion that they are stress granules. Five minutes after microinjection, these granules were approximately 0.50μm ± 0.22 μm in diameter, which is smaller than the typical stress granules observed after >30 minutes of oxidative stress, which measure around 1–2.0 μm (Anderson and Kedersha, 2009; Wheeler et al., 2016).

To examine how these granules change over time, we looked for colocalization of *ftz* mRNA with G3BP1 and TIA1 in individual RNA foci by fluorescence microscopy after 5, 10 and 15 minutes and analyzed the extent of colocalization by Pearson correlation analysis. The colocalization of *ftz* mRNA with both G3BP1 and TIA-1 in foci increased between 5 and 15 minutes after microinjection (**Fig. 1 E**). The granules grew larger until at 30 minutes they were 1.2μm ± 0.54 μm in diameter (for example, see **Fig. 1 F, S1 C**). Cells microinjected with *β-globin* mRNA also formed granules that concentrated *β-globin* mRNA and G3BP1 (**Fig. S1 D**), demonstrating that this phenomenon is not specific to *ftz* mRNA. By contrast, cells injected with FITC-dextran alone had no granules (**Fig. S1 E**), indicating that these structures form in response to the microinjected RNA. The granules that formed in response to microinjected *ftz* mRNA colocalized with endogenous *NORAD* lncRNA (**Fig. 1 G**), which is known to concentrate in stress granules, but not with endogenous *GAPDH* mRNA (**Fig. S1 F**), which does not (Khong et al., 2017). They were also seen by FISH with poly(A) mRNA (**Fig. S1 A**), suggesting that they accumulate a substantial fraction of all cytoplasmic mRNAs.

Together, these data demonstrate that *ftz* mRNA forms granules within 5 minutes after microinjection, which become larger and accumulate stress granule-specific RNA and proteins over the course of 30 minutes. Since these structures resembled stress granules in form, structure, and content, we will refer to them as stress granules, although they may differ from stress granules induced by oxidative, osmotic, or heat stresses in ways we have not assessed.

Fifteen minutes after microinjection, the *ftz* mRNA began to accumulate in another type of cytoplasmic structure: small foci that tended to be adjacent, or ‘docked’ to the sides of the stress granules containing *ftz* mRNA. After 30 minutes, the FISH signal from *ftz* mRNA in these foci was significantly greater than that in their adjacent stress granules (**Fig. S1 C**, green arrows). When compared with the stress granules containing G3BP1 and TIA-1, these foci were smaller, contained higher concentrations of *ftz* mRNA, and they colocalized with Dcp1a, a marker of processing (P)- bodies (**Fig. 1 F**). Although neighboring, uninjected cells (**Fig. 1 F**, yellow arrows) lacked granules containing TIA-1, they contained foci of Dcp1a, consistent with previous reports that P-bodies are constitutively present in most cells (Luo et al., 2018; Standart and Weil, 2018; Hubstenberger et al., 2017). All the granules that contained *ftz* mRNA but not TIA-1 also contained Dcp1a (**Fig. 1 F**); we will refer to these structures as P-bodies.

To examine how these P-bodies change over time, we looked for colocalization of *ftz* mRNA with TIA-1 or Dcp1a in individual RNA foci by fluorescence microscopy at intervals from 15 minutes to 4 hours after microinjection and analyzed the extent of colocalization by Pearson correlation analysis. By microscopy, we distinguished P-bodies that were docked to stress granules (see, for example, **Fig. S1 C**, green arrows) from those that were undocked (**Fig. S1 C,** magenta arrows). Fifteen minutes after microinjection, most P-bodies were docked to stress granules and contained significant amounts of TIA-1 and some Dcp1a (**Fig. 1 H**). Later, the P-bodies became more segregated from the stress granules and contained higher concentrations of Dcp1a and lower concentrations of TIA-1 than they did at earlier times (**Fig. 1 H**, movement of yellow and red puncta to top left quadrant over time). Four hours after microinjection, the two structures were largely distinct. We conclude from this analysis that Dcp1a-positive *ftz* mRNA granules gradually lose their stress granule identity (i.e. TIA-1 content) over time and become more distinct. This suggests the interesting possibility that mRNA is transferred from P-bodies to stress granules. Then, 30 minutes to 4 hours later, P-bodies containing *ftz* mRNA become dissociated from stress granules. Alternatively, docked P-bodies may dissolve over time to be replaced by undocked P-bodies containing *ftz* mRNA.

### Microinjected mRNA triggers stress granule formation by PKR-dependent and - independent mechanisms

Most forms of stress that induce stress granule formation – increased temperature, exposure to osmotic and oxidative stress, or infection with a virus, for example – inhibit the initiation of mRNA translation by a mechanism that depends on phosphorylation of the crucial translation initiation factor eIF2α by one of four stress-sensing kinases: PKR, the PKR-like endoplasmic reticulum kinase (PERK), heme-regulated inhibitor kinase (HRI), and GCN2 (Donnelly et al., 2013). This inhibition of mRNA translation, in turn, activates stress granule formation. A few stressors, however, trigger formation of stress granules independently of eIF2α phosphorylation (Dang et al., 2006; Kedersha et al., 2016; Aulas et al., 2017). To test whether microinjected mRNA induces stress granules with or without eIF2α phosphorylation, we quantified phospho-eIF2α levels by immunofluorescence microscopy. Microinjection of either *ftz* or *β-globin* mRNA significantly increased the intensity of phospho-eIF2α staining (**Fig. 2 A** and **S2 A,** quantified in **Fig. 2 B)**, when compared to uninjected cells or cells injected with buffer and the injection marker, FITC 70kDa dextran (**Fig. S2 B**).

**Figure 2.**
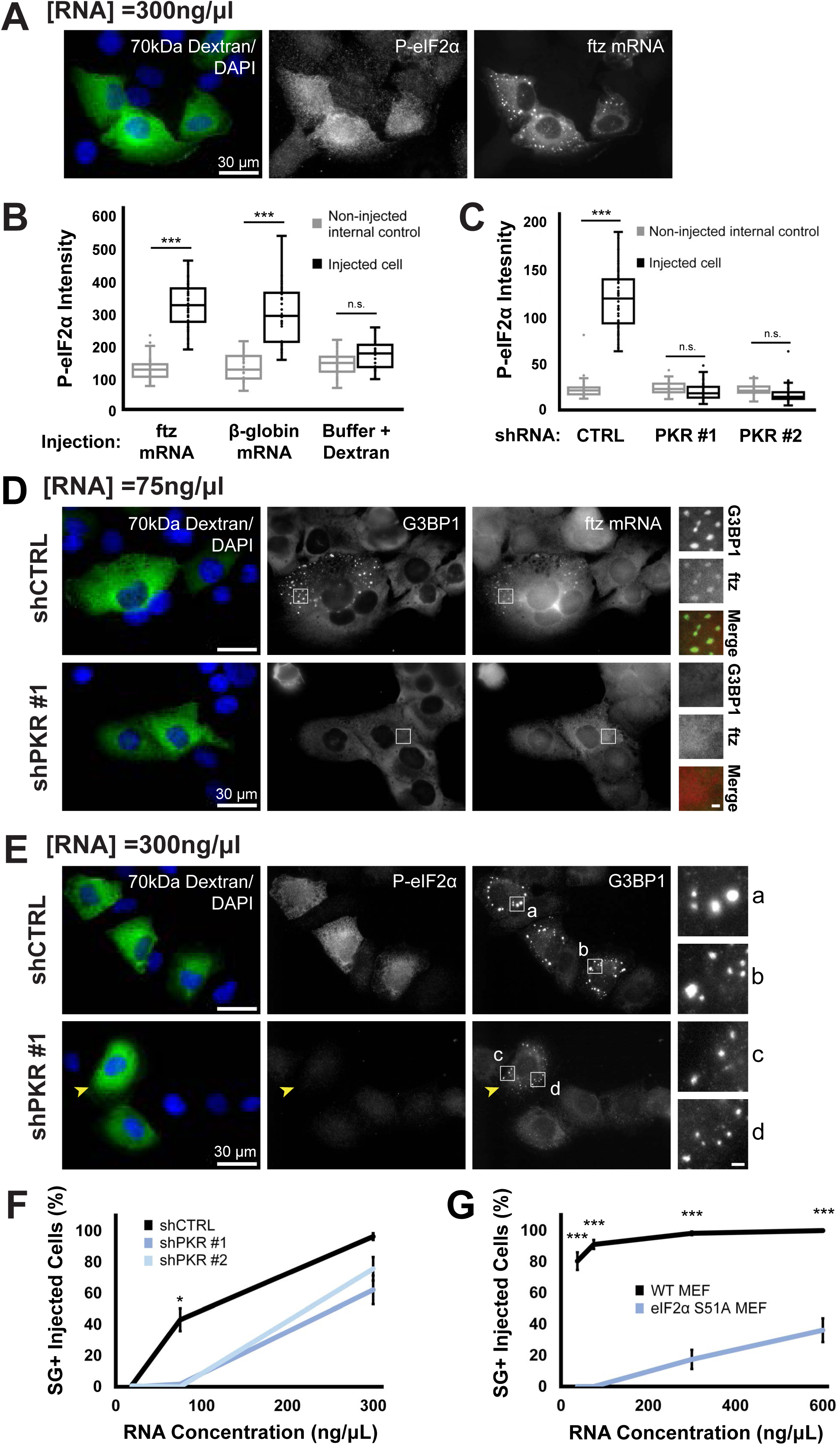
Microinjected mRNA triggers eIF2α phosphorylation via PKR, which is required for stress granule formation at low, but not high, concentrations of microinjected mRNA. **(A)** U2OS cells were microinjected with *ftz* mRNA (300ng/μL) and 70kD dextran conjugated to FITC. Cells were incubated for 30 minutes, fixed and stained for the injected mRNA using specific FISH probes, phospho-eIF2α by immunofluorescence, and DNA by DAPI. Scale bars: 30µm. **(B)** U2OS cells were microinjected, fixed, stained, and imaged as in (A). The total immunofluorescence intensity of phospho-eIF2α of these cells was quantified in arbitrary units (a.u.). Non-injected internal control cells are within the same field-of-view as injected cells. Each box plot displays the average, quartiles and standard deviations. (Not significant (n.s.): p>0.05; ***: p<0.001) **(C)** U2OS cells were transduced with lentivirus delivered shRNAs against PKR (two clones were used, PKR #1 and PKR #2) or a scrambled shRNA. Control or PKR-depleted U2OS cells were microinjected, fixed, stained, and imaged as in (E). The total immunofluorescence intensity of phospho-eIF2α of these cells was quantified in arbitrary units (a.u.). Non-injected internal control cells are within the same field-of-view as injected cells. Each box plot displays the average, quartiles and standard deviations. (Not significant (n.s.): p>0.05; ***: p<0.001) **(D)** Control or PKR-depleted U2OS cells were microinjected with *ftz* mRNA (75ng/μL) and 70kD dextran conjugated to FITC, fixed 30 minutes after injection, and stained with FISH probes against *ftz* mRNAs and immunostained against G3BP1. Scale bars: whole cell=30µm, insets=2µm. Zoomed in images are shown in boxed insets. The merged insets show G3BP1 in green and *ftz* mRNA in red. **(E)** Control or PKR-depleted U2OS cells were microinjected with *ftz* mRNA (300ng/μL) and 70kD dextran conjugated to FITC, fixed 30 minutes after injection, and immunostained against G3BP1 and phospho-eIF2α. A single PKR-depleted cell with stress granules are marked with an yellow arrow. Scale bars: whole cell=30µm, insets=2µm. Zoomed in images are shown in boxed insets. **(F)** Control or PKR-depleted U2OS cells were microinjected with various amounts of mRNA, fixed, stained, and imaged as in (D and E). The percentage of microinjected cells with visible SGs was quantified. Each data point represents the average and standard error of 3-6 experiments. (ns: p>0.05; *: p<0.05). **(G)** Wildtype or eIF2α-S51A mutant mouse embryonic fibroblasts (MEFs) were microinjected with various amounts of mRNA, fixed, stained and imaged as in (Supplementary Figure 2D-E). The percentage of microinjected cells with visible SGs was quantified. Each data point represents the average and standard error of 6 experiments. (ns: p>0.05; ***: p<0.001).

To determine if microinjected mRNA triggers PKR-mediated eIF2α-phosphorylation, we depleted PKR from U2OS cells by using a lentivirus vector to deliver shRNAs (**Fig. S2 C**), then assessed phospho-eIF2α levels and stress granule formation after *ftz* mRNA microinjection. This microinjected mRNA did not induce eIF2α phosphorylation in PKR-depleted cells (**Fig. 2 C**), consistent with previous finding using transfected mRNA (Kirschman et al., 2017). We found, however, that stress granule formation depended on the presence of PKR only when low concentrations of mRNA were injected. When the concentration of *ftz* mRNA microinjected was 75 ng/μl, it induced few or no stress granules in PKR-depleted cells, although it induced a significant number in ∼40% of control shRNA-treated cells (**Fig. 2 D**, quantified in **2 F**). By contrast, when the concentration of *ftz* mRNA microinjected was 300 ng/μl, it induced stress granules in about 60–70% of PKR-depleted cells (**Fig. 2 E**, quantified in **2 F**), demonstrating that high concentrations of microinjected mRNA trigger stress granules in PKR-independent manner. Consistent with this, in PKR-depleted cells injected with the higher concentration of mRNA we observed cells with stress granules that had no detectable phospho-eIF2α (**Fig. 2 E**, yellow arrow). These stress granules were smaller (0.80μm ± 0.43μm) than those in control shRNA-treated cells (1.3μm ± 0.57μm) (**Fig. 2 E**, compare insets a and b to c and d). These data indicate that microinjected mRNA triggers stress granule formation by a mechanism that involves PKR-mediated eIF2α phosphorylation at low concentrations of mRNA but that is independent of PKR and phospho-eIF2α at higher concentrations.

To confirm that high concentrations of microinjected mRNAs induce formation of stress granules independent of eIF2α phosphorylation, we assessed stress granule formation in mouse embryonic fibroblasts (MEFs) that were genetically altered to express only a mutant version of eIF2α (S51A) that cannot be phosphorylated by PKR (Scheuner et al., 2006; Dar et al., 2005).

Microinjection of low concentrations (75 ng/μl or less) of mRNA induced no stress granules in phospho-deficient eIF2α S51A MEFs, but strongly induced them in control cells, as expected (**Fig. 2 G**, representative images in **S2 D**). By contrast, microinjection of high concentrations (300 or 600 ng/μl) of mRNA induced stress granules in 20–40% of injected eIF2α S51A mutant MEFs (**Fig. 2 G**, representative images in **S2 E**). The stress granules induced in the eIF2α S51A mutant MEFs were smaller than those induced in wild-type MEFs, but they contained both *ftz* mRNA and G3BP1 (**Fig. S2 E**, see insets). Thus, high concentrations of microinjected mRNAs trigger the formation of stress granules without the need for phosphorylation of eIF2α on S51 by PKR, albeit less effectively than when eIF2α is phosphorylated.

We conclude from these experiments that mRNAs microinjected at low concentrations trigger formation of stress granules by a mechanism that requires PKR to phosphorylate eIF2α, whereas mRNAs microinjected at higher concentrations trigger formation of stress granules independently of both PKR and phospho-eIF2α.

### Pseudouridine in mRNA inhibits stress granule formation by the PKR–dependent mechanism

Messenger RNAs containing pseudouridine in place of uridine, when transfected into cells, activate PKR less than normal mRNAs do (Anderson et al., 2010). We investigated, therefore, whether microinjection of mRNAs containing pseudouridine would induce PKR-dependent stress granule formation. To do so, we transcribed *ftz* mRNA *in vitro* with pseudoUTP in place of UTP (to produce pseudouridine (Ψ)-*ftz* mRNA; see **Fig. S3 A**). We microinjected the Ψ-*ftz* mRNA or normal, uridine-containing *ftz* mRNA into U2OS cells and analyzed stress granule formation by immunofluorescence microscopy, as before. A low concentration (75 ng/μl) of microinjected Ψ-*ftz* mRNA induced no stress granules whereas the same concentration of normal *ftz* mRNA induced them in over 90% of injected cells (**Fig. 3 A**, quantified in **3 C**). Only when the concentration of microinjected Ψ-*ftz* mRNA was increased to >100 ng/μl, did we observe stress granules, and at 300 ng/μl we saw almost as many cells with stress granules as when we used 300 ng/μl of normal *ftz* mRNA (**Fig. 3 B**, quantified in **3 C**). We analyzed the amount of phospho-eIF2α in cells microinjected with 300 ng/μl of Ψ-*ftz* mRNA or normal *ftz* mRNA by immunofluorescence microscopy and observed that the cells microinjected with Ψ-*ftz* mRNA contained little phospho-eIF2α whereas those microinjected with the same quantity of normal *ftz* mRNA contained significantly more (**Fig. 3 B**). Quantification of the phospho-eIF2α fluorescence intensity showed that those microinjected with low or high concentrations of Ψ-*ftz* mRNA contained similar, little amounts of phospho-eIF2α as non-injected cells, whereas cells microinjected with normal *ftz* mRNA showed a concentration-dependent increase in phospho-eIF2α (**Fig. 3 D**). These observations indicate that pseudouridine-containing mRNAs do not induce eIF2α - phosphorylation, likely because they do not activate PKR and thus they do not induce the PKR-dependent mechanism of stress granule formation. High concentrations of pseudouridine-containing mRNAs, however, induce stress granules, further confirming that they act by a PKR-independent mechanism.

**Figure 3.**
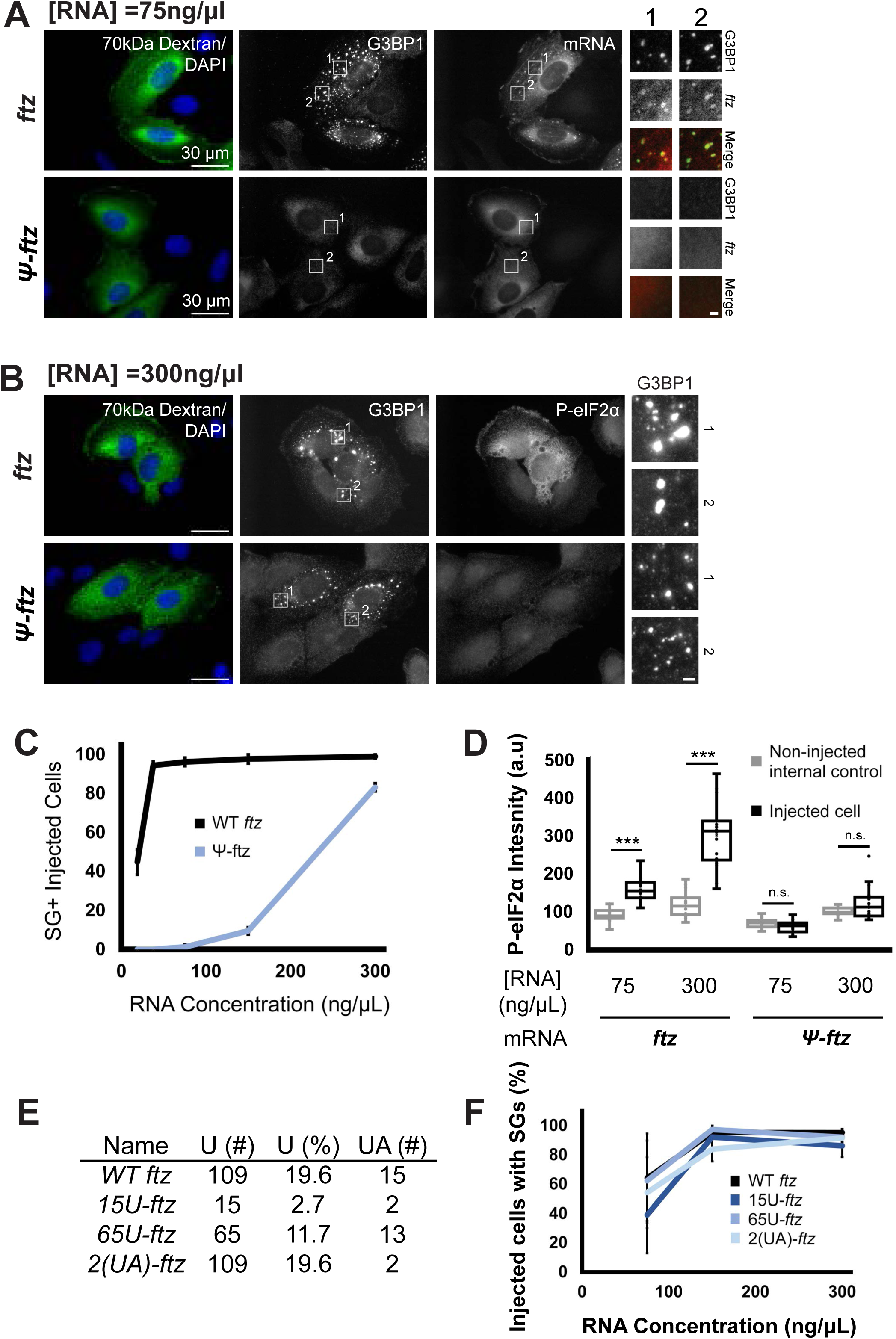
Pseudouridine incorporation in mRNA, but not the absence of uridine or UpA, block stress granule formation at low, but not high, concentrations. **(A-B)** U2OS cells were microinjected with either uridine-containing ftz mRNA or pseudouridine-containing *ftz* mRNA (“ψ-*ftz*”) at 75ng/μL (A) or 300ng/μL (B) and 70kD dextran conjugated to FITC. After 30 minutes, cells were fixed and stained for the SG-marker G3BP1 (A-B) and phospho-eIF2α (A) by immunofluorescence, *ftz* mRNA using FISH probes (B), and DNA by DAPI. Scale bars: whole cell=30µm, insets=2µm. Zoomed in images are shown in boxed insets. The merged insets show G3BP1 in green and *ftz* mRNA in red. **(C)** U2OS cells were microinjected with 18.75, 37.5, 75, 150 or 300ng/μL of *ftz* mRNA, fixed, stained, and imaged as in (B). The percentage of microinjected cells with visible SGs was quantified. Each data point represents the average and standard error of 3 experiments. (**: p<0.01; ***: p<0.001). **(D)** U2OS cells were microinjected with 75ng/μL or 300ng/μL of mRNA, fixed, stained, and imaged as in (A-B). The total immunofluorescence intensity of phospho-eIF2α of these cells was quantified in arbitrary units (a.u.). Non-injected internal control cells are within the same field-of-view as injected cells. Each box plot displays the average, quartiles and standard deviations. (Not significant (n.s.): p>0.05; ***: p<0.001) **(E)** A table showing *ftz* mRNA constructs with varying uridine content by the number and percentage of uridines in the transcripts, and the number of UA dinucleotides. **(F)** U2OS cells were microinjected with WT-, 15U-, 65U-, or 2(UA)-*ftz* mRNA at various various concentrations (75, 150 or 300ng/μL), fixed, stained, and imaged as in (Supplementary Figure 3C-D). The percentage of microinjected cells with visible SGs was quantified. Each data point represents the average and standard error of 3 experiments. (ns: p>0.05).

The failure of low concentrations of Ψ-*ftz* mRNA to induce stress granules begs the question of whether it is the presence of the pseudouridine or the lack of normal uridine in the *ftz* mRNA that causes this effect. To answer this question, we synthesized versions of *ftz* mRNA containing various amounts of uridine (**Fig. 3 E, S3 B**) by generating three mutant mRNAs: *15U-ftz*, in which 94 of the 109 uridines in *ftz* mRNA were mutated; *65U-ftz* in which 44 uridines were mutated, and *2(UA)ftz* in which 13 of the 15 UpA dinucleotides in *ftz* mRNA were mutated. (UpA dinucleotides are generally under-represented in virus RNAs (Mordstein et al., 2021), potentially to evade PKR-mediated immune responses.) Each mutant was modified to maintain a similar total free energy (**Table S1,** (Zuker, 2003)). All three mutant versions of *ftz* mRNA stimulated stress granule formation as robustly as the original *ftz* mRNA (**Fig. 3 F,** representative images in **S3 C and D**), indicating that changing the amount of uridine or UpA dinucleotides does not influence stress granule formation. These findings demonstrate that it is the presence of pseudouridine, rather than the absence of uridines or UpA dinucleotides, in the microinjected mRNA that prevents stress granule formation by the PKR-dependent mechanism.

### dsRNA is required for stress granule formation by the PKR–dependent mechanism

How might the presence of pseudouridine in microinjected mRNA prevent stress granule formation by the PKR-dependent mechanism? The T7 RNA polymerase we used to synthesize the mRNAs was reported to produce a small amount of dsRNA by antisense transcription from the DNA template and/or transcription off of the newly synthesized RNA (Cazenave and Uhlenbeck, 1994; Mu et al., 2018; Wu et al., 2020). The presence of certain nucleotide analogues, including pseudouridine, in the T7 RNA polymerase reaction inhibit formation of these dsRNA byproducts (Mu et al., 2018). We hypothesized, then, that pseudouridine might block stress granule formation by inhibiting the formation of dsRNA byproducts by T7 RNA polymerase.

To test this hypothesis, we treated the *in vitro* transcription products of T7 RNA polymerase with RNase III, which cleaves dsRNA (Court et al., 2013), and then purified the *ftz* mRNA. The untreated and RNase III-treated mRNAs were analyzed on a 1% agarose gel. By loading a large amount of mRNA (4 µg per lane), we could detect bands of an apparently higher molecular weight than the majority of mRNA (**Fig. 4 A**, RNaseIII - panel red asterisks); after RNase III treatment, however, we no longer observed these higher molecular weight bands (**Fig. 4 A**, RNaseIII + panel). We microinjected high and low concentrations of the RNase III-treated or untreated *ftz* mRNA into U2OS cells and analyzed stress granule formation by fluorescence microscopy, as before. The high concentration of RNase III-treated *ftz* mRNA (300 ng/μl) induced stress granules (**Fig. 4 B**), whereas the low concentration (75 ng/μl) did not (**Fig. 4 C**). By contrast, both concentrations of untreated *ftz* mRNA induced stress granules (**Fig. 4 B** and **4 C**). Quantification of the number of cells containing stress granules showed that the high concentration of RNase III-treated *ftz* mRNA induced stress granules in about half as many cells as did the high concentration of untreated *ftz* mRNA (**Fig. 4 D**). Together, these data suggest that stress granule formation by the PKR-dependent mechanism at low concentrations of microinjected mRNA is driven by dsRNA, whereas much of stress granule formation by the PKR-independent mechanism at high concentrations of microinjected mRNA is not.

**Figure 4.**
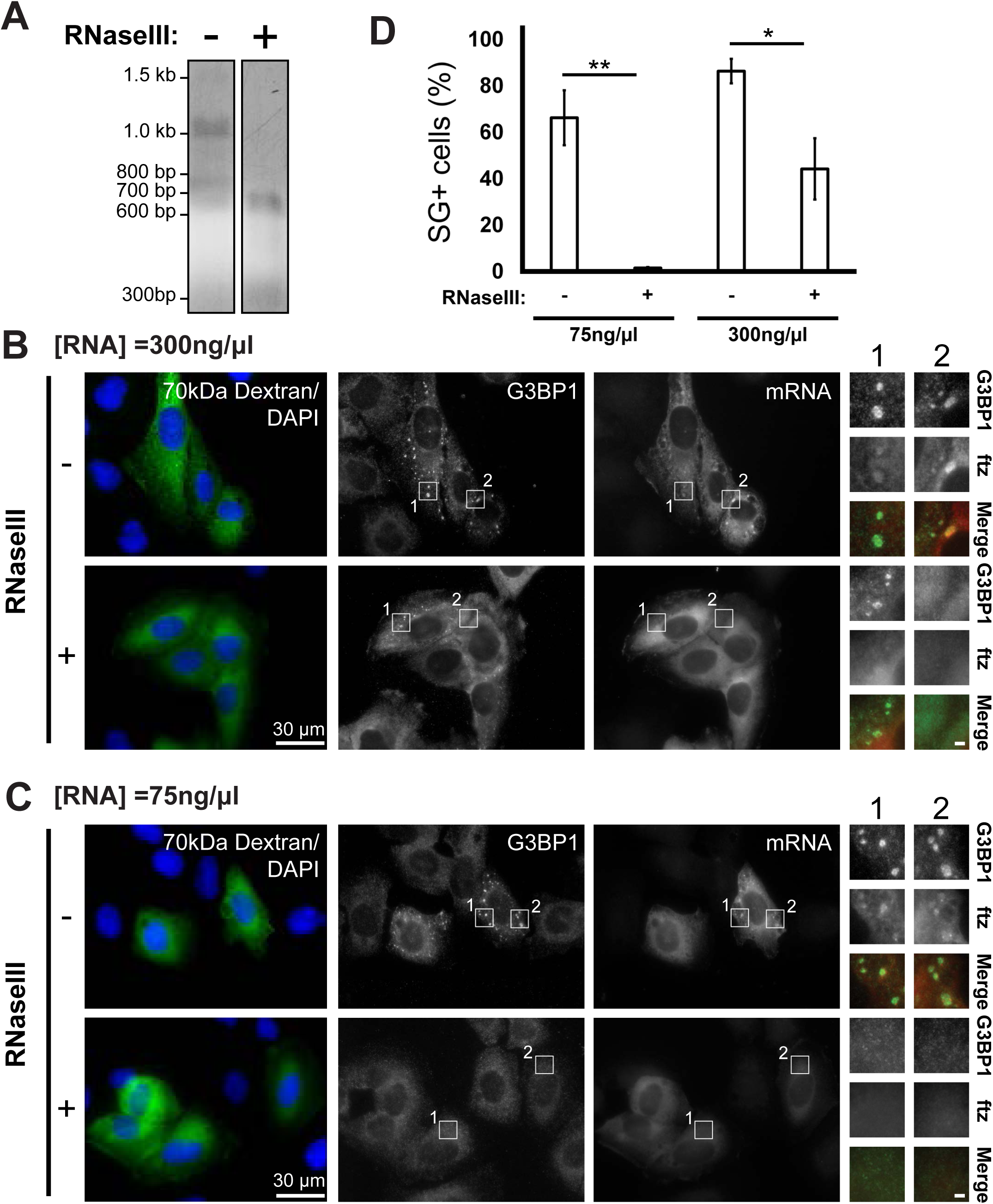
RNaseIII treated mRNA block stress granule formation at low, but not high, concentrations. **(A)** *ftz* mRNAs were *in vitro* transcribed, capped and polyadenylated, then purified. Samples untreated (“-“) or RNaseIII-treated (“+”) were collected and resolved on an a 1% agarose gel. The migration of standards are indicated on the right. **(B-C)** U2OS cells were microinjected with either untreated or RNaseIII-treated *ftz* mRNA at 300ng/μL (B) or 75ng/μL (C) and 70kD dextran conjugated to FITC. After 30 minutes, cells were fixed and stained for the SG-marker G3BP1 by immunofluorescence, *ftz* mRNA using FISH probes and DNA by DAPI. Scale bars: whole cell=30µm, insets=2µm. Zoomed in images are shown in boxed insets. The merged insets show G3BP1 in green and *ftz* mRNA in red. **(D)** The percentage of microinjected cells with visible SGs in cells imaged in (B-C) were quantified. Each column and error bar represent the average and standard error of 3 experiments. *: p<0.05; **: p<0.01).

### G3BP1 and G3BP2 enhance stress granule formation by both mechanisms

To understand better the mechanisms driving PKR-dependent and -independent stress granule formation, we investigated the contribution of the stress granule scaffolding proteins G3BP1 and G3BP2, which are key mediators of stress granule formation in response to various forms of cell stress (Matsuki et al., 2013; Kedersha et al., 2016; Guillén-Boixet et al., 2020; Yang et al., 2020). We microinjected low and high concentrations of *ftz* mRNA into wild-type U2OS cells and G3BP1–G3BP2 double deletion (ΔΔG3BP1/2) U2OS cells, which were generated previously by CRISPR/Cas9 gene editing (Kedersha et al., 2016). We visualized stress granules by immunostaining FMRP, another stress granule component. Microinjection of a low concentration of *ftz* mRNA (75 ng/μl) into ΔΔG3BP1/2 U2OS cells failed to induce stress granules, whereas it induced granules in many wild-type cells, as expected (**Fig. S4 A**, see quantifications in **Fig. 5 A**). Thus, we conclude that G3BP1 and/or G3BP2 are required for stress granule formation by the PKR-dependent mechanism. Microinjected mRNA triggered eIF2α phosphorylation in both ΔΔG3BP1/2 and control U2OS cells by immunofluorescence (**Fig. S4 B,** see quantifications in **Fig. 5 B**), indicating that G3BP1/2 likely act downstream of PKR and eIF2α phosphorylation in triggering stress granule formation. Consistent with this, some microinjected ΔΔG3BP1/2 cells were positive for phospho-eIF2α but lacked stress granules (**Fig. S4 C**).

**Figure 5.**
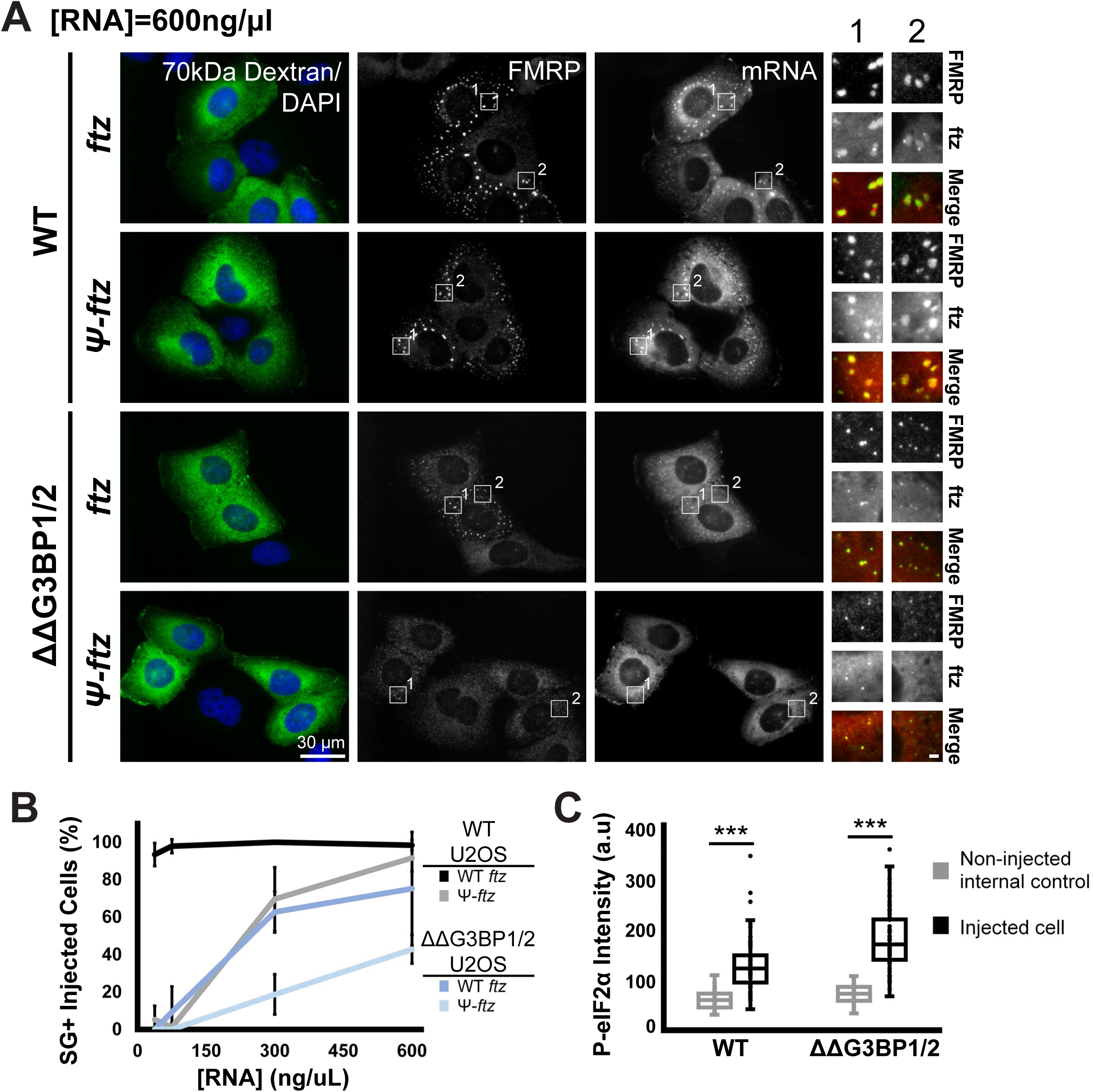
G3BP1 and G3BP2 enhance stress granule formation downstream of PKR and by PKR-independent mechanism. **(A)** WT and ΔΔG3BP1/2 U2OS cells were microinjected with 37.5, 75, 300 or 600 of mRNA, fixed, stained, and imaged as in (A). The percentage of microinjected cells with visible SGs was quantified. Each data point represents the average and standard deviation of 2 experiments. **(B)** WT and ΔΔG3BP1/2 U2OS cells were microinjected, fixed, stained for phospho-eIF2α, and imaged as in (Supplemental Figure 4A). The total immunofluorescence intensity of phospho-eIF2α of these cells was quantified in arbitrary units (a.u.). Non-injected internal control cells are within the same field-of-view as injected cells. Each box plot displays the average, quartiles and standard deviations. (Not significant (n.s.): p>0.05; ***: p<0.001). **(C)** WT and ΔΔG3BP1/2 U2OS cells were microinjected with uridine-containing (“U-*ftz*”) or pseudouridine-containing *ftz* mRNA (“ψ-*<u>ftz</u>*”) at 600ng/μL and 70kD dextran conjugated to FITC. After 30 minutes, cells were fixed and stained for ftz mRNA using FISH probes, the SG-marker FMRP by immunofluorescence, and DNA by DAPI. Scale bars: whole cell=30µm, insets=2µm. Zoomed in images are shown in boxed insets. The merged insets show G3BP1 in green and *ftz* mRNA in red.

Microinjection of a high concentration of *ftz* mRNA (300 ng/μl) induced stress granules in fewer ΔΔG3BP1/2 U2OS cells than it did in wild-type cells (see quantifications in **Fig. 5 A**): only 60–70% of ΔΔG3BP1/2 cells formed stress granules, whereas nearly all of the wild-type cells formed them. This suggests that G3BP1/2 play important roles in RNA-induced stress granule formation independent of PKR. To investigate this further, we injected Ψ-*ftz* mRNA, which does not activate PKR, into wild-type and ΔΔG3BP1/2 U2OS cells and again observed stress granule formation by immunofluorescence microscopy of FMRP. As expected from our findings in Figure 4, when we microinjected a low concentration of Ψ-*ftz* mRNA (75 ng/µl), no stress granules formed in either cell line (**Fig. S4 A)**. A high concentration of Ψ-*ftz* mRNA (600 ng/µl), by contrast, did induce stress granules, although significantly fewer in ΔΔG3BP1/2 cells than in wild-type U2OS cells (**Fig. 5 C;** see quantifications in **Fig. 5 B**). This finding is consistent with our observations with normal *ftz* mRNA and demonstrates that G3BP1 and/or G3BP2 are/is crucial for stress granule formation in response to microinjected mRNA regardless of its concentration.

We note that high concentrations of Ψ-*ftz* mRNA induced stress granules in 20–40% of ΔΔG3BP1/2 cells, indicating that mRNA can drive stress granule formation independent of PKR– phospho-eIF2α and G3BP1/2. Thus, G3BP1 and/or G3BP2 are not strictly required, but significantly enhance mRNA-induced stress granule formation.

### Stress granule formation requires acute increase of ribosome-free cytosolic mRNAs

How might a high concentration of mRNA microinjected into the cell cytoplasm induce stress granule formation by a mechanism that does not require PKR and phospho-eIF2α, or the presence of G3BP1 and G3BP2? One possible explanation is that the high concentration of protein-free mRNA microinjected into the cells engage in weak multivalent interactions with other RNAs and phase separate, bringing in RNA binding proteins. If so, simply releasing mRNA from ribosomes in the cytosol might be sufficient to drive stress granule formation. To test this possibility, we treated cells with inhibitors of translation that release mRNA from ribosomes, puromycin or homoharingtonine. Puromycin treatment of U2OS cells did not activate PKR (as indicated by the amount of phospho-T466-PKR) or induce eIF2α phosphorylation (**Fig. 6 A;** poly(I:C), a well-known activator of PKR (Tu et al., 2012) was used as a positive control), indicating that this inhibitor, and probably homoharingtonine also, acts downstream of PKR and phospho-eIF2α. Also, when compared to treatment of the cells with sodium arsenite, which causes oxidative stress, these two inhibitors induced stress granules in very few cells (**Fig. 6 B**), consistent with previous studies (Bounedjah et al., 2014; Kedersha et al., 2000). This indicates that simply releasing endogenous mRNA from ribosomes is not sufficient to promote stress granule formation by the PKR-independent mechanism.

**Figure 6.**
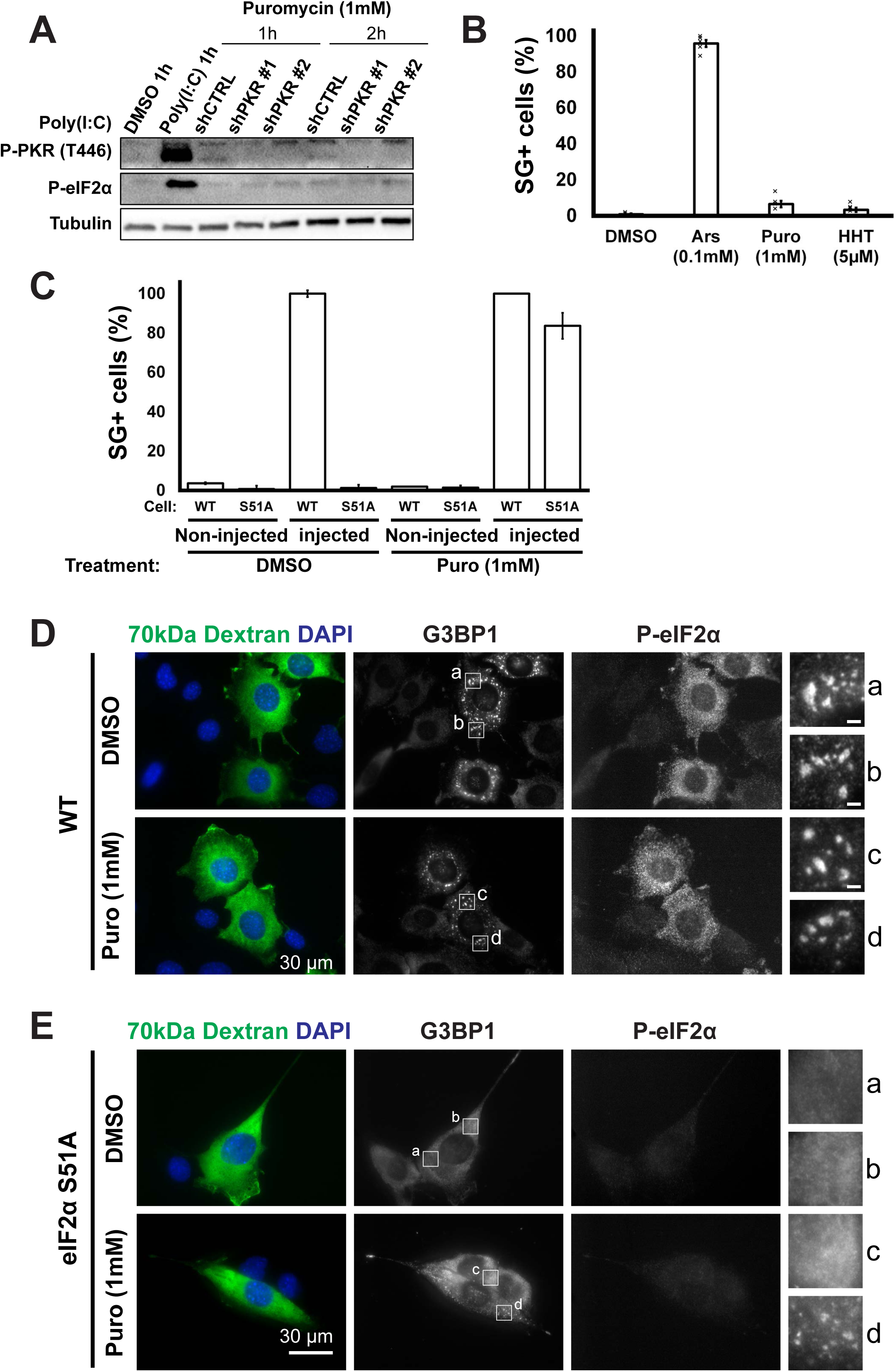
Stress granule formation requires tipping the balance between mRNA and RNA-binding proteins. **(A)** Control- or PKR-depleted U2OS cells were treated with puromycin (1µM) for 1 or 2 hours, and subsequently lysed, separated by SDS-PAGE and immunostained against phospho-PKR (Thr446), phospho-eIF2α and α-tubulin. Poly(I:C) transfection is a canonical inducer of PKR activation and phospho-eIF2α. The migration of standards are indicated on the right. **(B)** WT U2OS cells were treated with DMSO, sodium arsenite (“Ars,” 0.1mM), puromycin (“Puro,” 1mM), or homoharringtonine (“HHT,” 5µM) for 30 minutes, then fixed and stained for the SG-marker G3BP1 by immunofluorescence, and DNA by DAPI. The percentage of cells with visible SGs was quantified. Each column represents the average and standard deviation of 3-5 experiments. **(C)** Wild-type or eIF2α S51A phosphodeficient MEF cells were microinjected *ftz* mRNA (300ng/μL) and 70kD dextran conjugated to FITC, treated with DMSO or puromycin (1mM) for 30 minutes, fixed and stained against the SG-marker, G3BP1 (as shown in B and C). The percentage of cells with visible SGs in injected and non-injected cells were quantified. Each column represents the average and standard deviation of 3 experiments. **(D-E)** Wild-type (D) or eIF2α S51A phosphodeficient (E) MEF cells were microinjected *ftz* mRNA (300ng/μL) and 70kD dextran conjugated to FITC, treated with DMSO or puromycin (1mM) for 30 minutes, fixed and stained against *ftz* mRNA by FISH, G3BP1 by immunofluorescence, and DNA by DAPI. Scale bars: whole cell=30µm, insets=2µm. Zoomed in images are shown in boxed insets.

Although puromycin treatment releases most mRNAs from ribosomes, a significant portion of these ‘naked’ mRNAs will be bound quickly by RNA-binding proteins, reducing the effective concentration of naked mRNAs that can engage in RNA–RNA interactions in the cytosol. If so, acutely increasing the level of ribosome-free RNA by microinjection should saturate these RNA-binding proteins and drive stress granule formation upon puromycin treatment. To test this prediction in a PKR-independent manner, we microinjected 300 ng/µl *ftz* mRNA into MEFs that express only the S51A mutant of eIF2α that cannot be phosphorylated by PKR, and treated them with puromycin or with the solvent, DMSO, as a control. More than 80% of the microinjected cells treated with puromycin formed stress granules, a percentage similar to wild-type MEFs, whereas very few of those treated with DMSO did (**Fig. 6 C**), as in the experiment shown in Figure 2 G. Also, treating these cells with puromycin alone (i.e. without microinjecting mRNA) induced few or no stress granules in both wild-type and eIF2α S51A MEFs (**Fig. 6D and E**, see non-injected internal control in **Fig. 6 C**), as in WT U2OS cells. This suggests that stress granule formation by PKR-independent mechanism requires amounts of ribosome-free mRNA in the cytosol greater than those induced simply by releasing the endogenous ribosome-bound RNA.

## Discussion

Cytoplasmic ribosome-free mRNA, generated from global translational arrest and polysome disassembly, has been proposed to drive stress granule formation (Van Treeck et al., 2018; Van Treeck and Parker, 2018; Tian et al., 2020). In this study, we set out to test this hypothesis by using mRNA microinjection as a way to introduce ribosome-free mRNA in a controlled manner. We find that when microinjected into the cytoplasm, mRNAs transcribed *in vitro* by T7 RNA polymerase induces stress granules by two mechanisms: one dependent on PKR phosphorylation of eIF2α and, the other independent of PKR and phospho-eIF2α. Both mechanisms are promoted by the RNA-binding proteins G3BP1 and G3BP2, which have a scaffolding function in stress granules. Microinjected mRNA can induce stress granules independently of PKR, phospho-eIF2α, and G3BP1/G3BP2 only when the concentration of cytoplasmic mRNAs is greater than the endogenous concentration.

Our data indicate that even low concentrations of microinjected mRNA triggers stress granule assembly by the PKR-dependent mechanism. This is consistent with a previous finding that lipid-based transfection of cells with *in vitro* transcribed mRNAs activates PKR (Anderson et al., 2010; Kirschman et al., 2017), although how exogenous mRNA activates PKR was unclear until now. PKR is activated by dsRNA of 30 base pairs or longer (Lemaire et al., 2008), which induces PKR dimerization and activation (Gal-Ben-Ari et al., 2018). The exogenous RNAs that we used here and previously (Mahadevan et al., 2013) are not predicted to form dsRNA stretches longer than 30 base pairs. Why then do these mRNAs trigger PKR activation? We demonstrate here that PKR activation is blocked by including pseudouridine in the exogenous RNAs. One possible explanation for this observation is that our RNAs containing pseudouridine do form dsRNA but they are a poor substrates for PKR, as previously proposed (Anderson et al., 2010; Karikó et al., 2005). Another possibility is that the presence of pseudouridine in the *in vitro* transcription reaction inhibits the production of anti-sense transcripts generated by T7 RNA polymerase (Mu et al., 2018). The fact that RNase III treatment phenocopies pseudouridine treatment in our study is consistent with this latter possibility: PKR activation by exogenous mRNA is due to dsRNA byproducts of T7 polymerase, and pseudouridine blocks dsRNA generation. Production of dsRNA byproducts by T7 polymerase varies depending on the template sequence and reaction conditions (Mu et al., 2018; Wu et al., 2020). Our findings raise an important awareness that the findings of previously studies that examined PKR activation and stress granule formation using exogenous mRNA (Mahadevan et al., 2013; Bounedjah et al., 2014; Fay et al., 2017; Kirschman et al., 2017) may be confounded by the presence of trace amounts of dsRNA in the samples. Likewise, dsRNA contamination could influence interpretation of many other studies using *in vitro* transcribed mRNAs by T7 polymerase in cells and *in vitro*; it will be important to include additional steps to degrade or prevent dsRNA byproducts.

In the PKR-independent mechanism, microinjected mRNA triggered stress granule formation without PKR activity, eIF2 α phosphorylation, or G3BP1/2. One possibility is that excess protein-free mRNA stimulates stress granule condensation directly through transient RNA– RNA interactions of stretches shorter than the 30 base pairs of dsRNA required to activate PKR (Lemaire et al., 2008), as we do not detect eIF2α phosphorylation in these cells. This possibility is in line with previous proposals that RNA–RNA interactions drive stress granule formation (Van Treeck et al., 2018; Van Treeck and Parker, 2018; Tian et al., 2020). In contrast to these previous proposals, however, our data indicate that RNA–RNA interactions arising from polysome disassembly alone is insufficient to trigger stress granule formation; instead, stress granule formation requires additional protein-free mRNAs introduced by microinjection. We hypothesize that microinjection acutely increases the cytoplasmic pool of free mRNA to a concentration that saturates the capacity of the RNA-binding proteins to bind and prevent RNA–RNA interactions. These findings are also consistent with previous reports that certain compounds that disassemble polysomes are not sufficient to trigger stress granule formation alone, but do enhance stress granule formation in the presence of other triggers, such as arsenite-induced oxidative stress (Kedersha et al., 2000). An alternative explanation is that the excess microinjected mRNA activates some other RNA sensor, such as RIG-I or MDA5, by another mechanism (Corbet et al., 2023).

Interestingly, both the PKR-dependent and -independent mechanisms of stress granule formation were enhanced by the presence of the scaffolding proteins G3BP1 and G3BP2, indicating that formation of stress granules requires not only RNA–RNA interactions but also RNA–protein and likely protein–protein interactions. This is consistent with previous reports that the interaction of G3BP1 with RNA caused it to condense *in vitro* (Yang et al., 2020; Sanders et al., 2020; Guillén-Boixet et al., 2020), organizing RNA into much larger, rounder, and more dynamic condensates. Furthermore, RNA partitioning into stress granules is dictated by the synergistic effects of both RNA–RNA-binding protein and RNA–RNA interactions (Matheny et al., 2021). Together, these results suggest that while ribosome-free mRNA can drive stress granule formation, it is likely not sufficient to do so without perturbations that tip the balance of mRNA and RNA-binding protein interactions. One such condition might be viral infection, which introduces additional exogenous RNAs to the cytosol or promote the cleavage of G3BP1/2. Overall, our findings highlight the importance of RNA-binding proteins G3BP1 and G3BP2 in mRNA-driven stress granule formation, which likely applies to other stress conditions as well.

Future studies of how exogenous RNAs trigger stress granules may help us to uncover additional sensors of foreign RNAs that coordinate the innate immune response to infection. Such additional sensors might be targeted to diminish inflammation or to boost the translation of therapeutic mRNAs, such as those in mRNA-based vaccines. The presence of either pseudouridine or N1-methylpseudouridine drastically inhibits the efficiency of mRNA translation (Eyler et al., 2019). It would be interesting to dissect the exact effects of these nucleotide analogues on stress granule formation and stimulation of the innate immune response, as pseudouridine-free mRNA vaccines may have enhanced efficacy.

---

**Supplemental Table 1.**
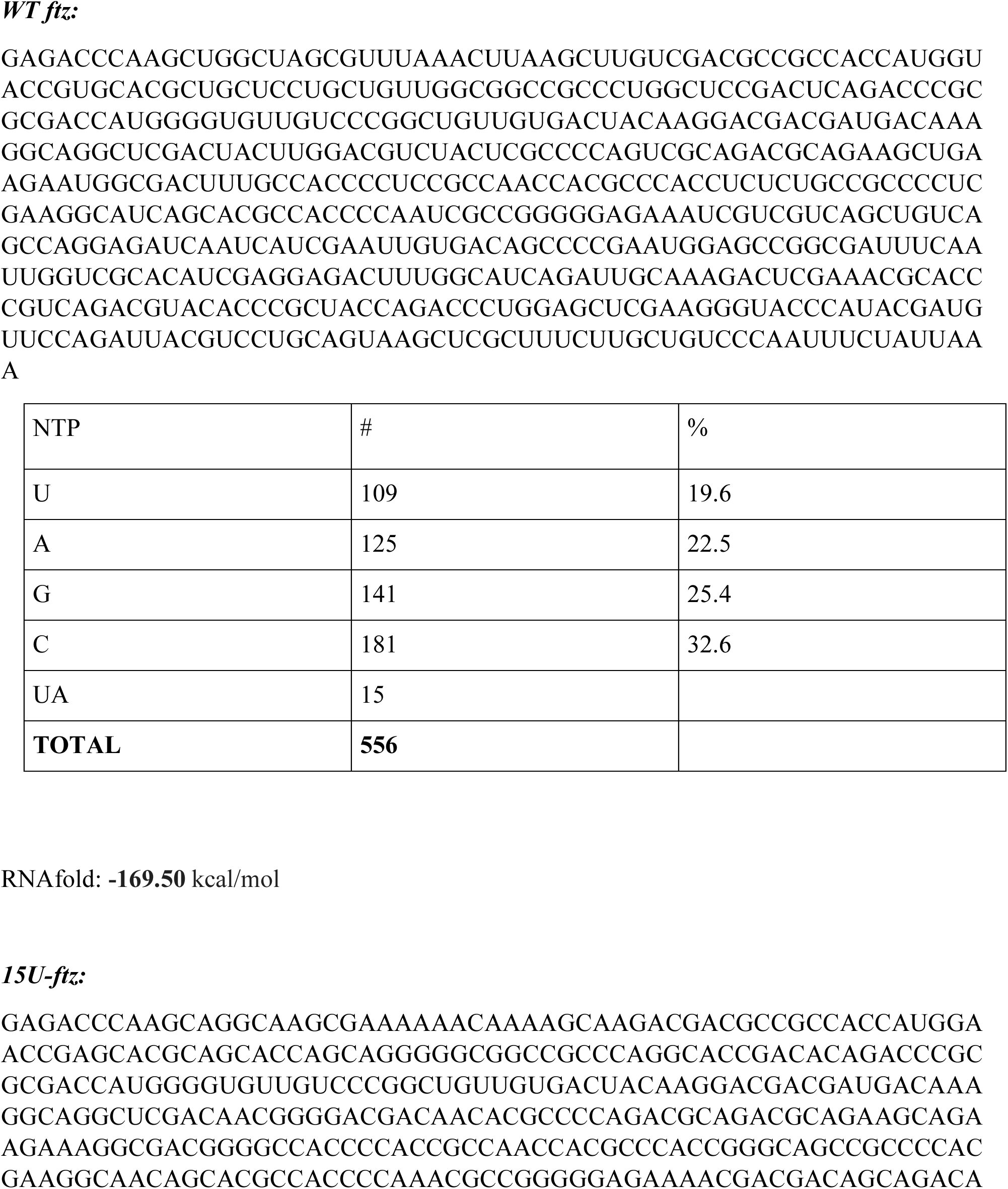
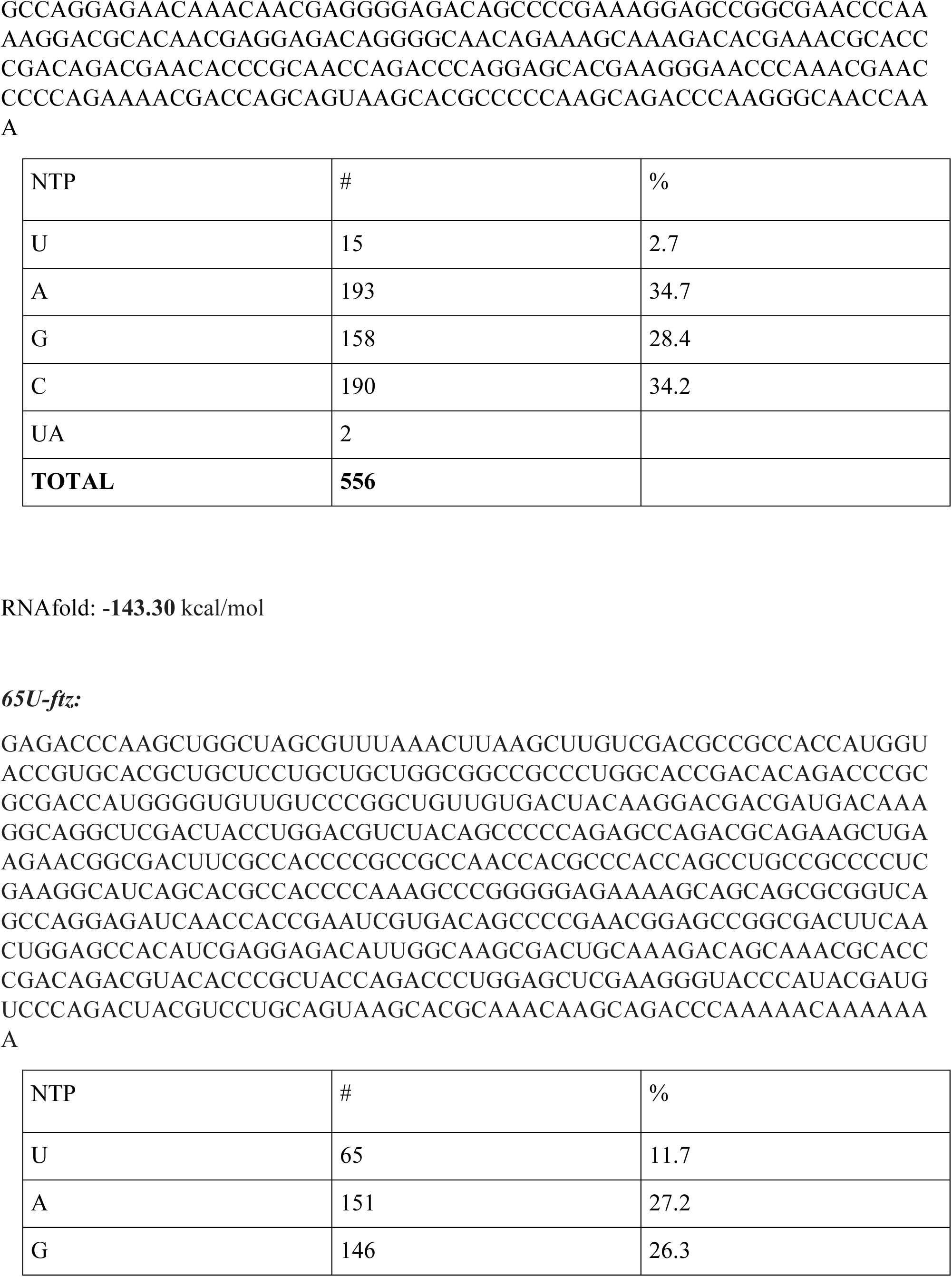
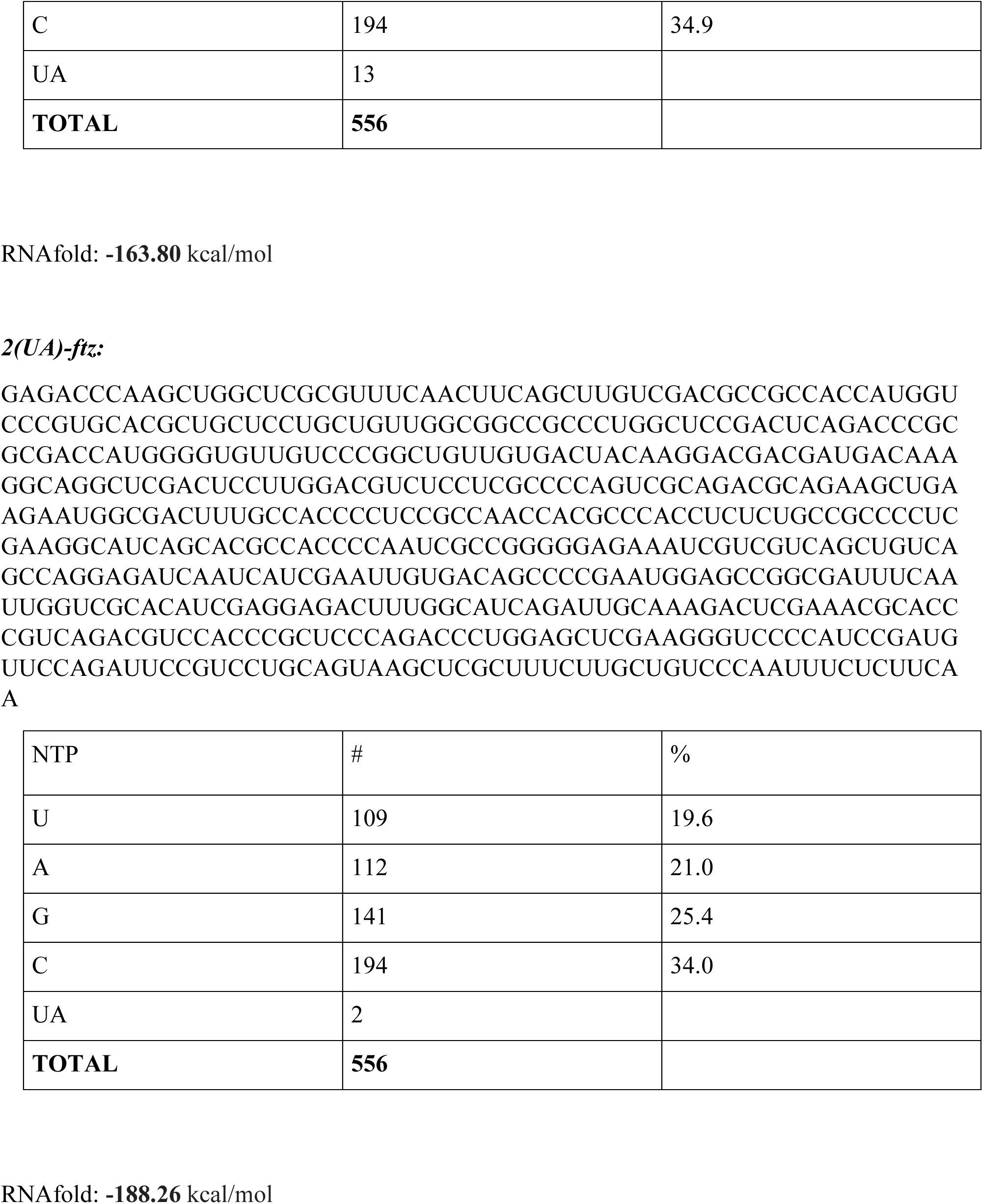
Sequence and features of *ftz* mRNA uridine and UpA mutants used in Figure 3.

| NTP | # | % |
| --- | --- | --- |
| U | 109 | 19.6 |
| A | 125 | 22.5 |
| G | 141 | 25.4 |
| C | 181 | 32.6 |
| UA | 15 |  |
| **TOTAL** | **556** |  |
| U | 15 | 2.7 |
| A | 193 | 34.7 |
| G | 158 | 28.4 |
| C | 190 | 34.2 |
| UA | 2 |  |
| **TOTAL** | **556** |  |

| NTP | # | % |
| --- | --- | --- |
| U | 65 | 11.7 |
| A | 151 | 27.2 |
| G | 146 | 26.3 |
| C | 194 | 34.9 |
| UA | 13 |  |
| **TOTAL** | **556** |  |

| NTP | # | % |
| --- | --- | --- |
| U | 109 | 19.6 |
| A | 112 | 21.0 |
| G | 141 | 25.4 |
| C | 194 | 34.0 |
| UA | 2 |  |
| **TOTAL** | **556** |  |

## Materials and methods

### Plasmids constructs and primers

The MHC-*ftz-Δi* and *βG-Δi* reporter constructs in pcDNA3.0 were previously described (Palazzo et al., 2007; Akef et al., 2013). The MHC-*ftz-Δi* sequences with varying uridine-content as detailed in Supplemental Table 1 were synthesized in pJET1.2 by General Biosystems.

### Antibodies

Antibodies used in this study were PKR (Abcam, ab32052), P-PKR (T446) (Abcam, ab32036), P-eIF2α (Cell Signaling Technology, #9721), G3BP1 (Abcam, ab56574), Caprin (Proteintech, 15112-1-AP), TIA-1 (Invitrogen, MA5-26474), FMRP (Cell Signaling Technology, #4317), PABP (Abcam, ab21060) Dcp1a (Sigma, WH0053802M6), and tubulin (Sigma, T6199).

### Cell culture and lentiviral-mediated shRNA protein depletion

Human osteosarcoma cells (U2OS), embryonic kidney 293T cells (HEK293T), and mouse embryonic fibroblasts (MEF) were grown in DMEM media (Wisent) supplemented with 10% fetal bovine serum (FBS) (Wisent) and 5% penicillin/streptomycin (Wisent). The lentiviral delivered shRNA protein depletion was performed as previously described (Akef et al., 2013; Lee et al., 2020). Briefly, HEK293T were plated at 40% confluency on 60 mm dishes and transfected with the gene specific shRNA pLKO.1 plasmid (Sigma), packaging plasmid (Δ8.9) and envelope (VSVG) vectors using Lipo293T DNA *in vitro* transfection reagent (SignaGen) according to the manufacture’s protocol. Viruses were harvested from the media 48 hours later and added to U2OS cells pre-treated with 8 μg/ml hexadimethrine bromide. U2OS cells were then selected with 2 μg/ml puromycin media for at least 4 to 6 days. Western blotting was used to determine the efficiency of PKR depletion. The shRNAs constructs (Sigma) used in this study are as follows:

‘PKR#1’shRNA

TRCN0000197012(CCGGGAGGCGAGAAACTAGACAAAGCTCGAGCTTTGTCTAGTTTCTCGCCTCTTTTTTG).

‘PKR#2’shRNA,

TRCN0000196400(CCGGGCTGAACTTCTTCATGTATGTCTCGAGACATACATGAAGAAGTTCAGCTTTTTTG).

### *In vitro* mRNA synthesis

MHC-ftz-Δi mRNA was synthesized *in vitro* using HiScribe T7 High Yield RNA Synthesis Kit (New England Biolabs) with MHC-ftz-Δi in pcDNA3.0 (30) digested with XhoI as a template. Template DNA was subsequently removed from the mRNA synthesis reaction with DNase I incubation at 37°C for 20 minutes. The RNA was capped with Vaccinia Capping System (New England Biolabs) and polyadenylated with E. coli Poly(A) Polymerase (New England Biolabs), following the manufacturers’ protocol. For pseudouridine-incorporated mRNA, all of the UTP in the reaction was replaced by Ψ (TriLink). The *in vitro* synthesized mRNA was purified, evaluated via spectroscopy, and resolved on an agarose gel (%1).

### Microinjection

For microinjection experiments, cells were plated on 22 × 22 mm coverslips (VWR) in 35 mm mammalian tissue culture dishes (Thermo Scientific) for 24 h prior to injection. Microinjections were performed as previously described (Gueroussov et al., 2010). RNA was microinjected at indicated concentrations with 70 kDa FITC Green Dextran (1/5^th^ volume) in Injection Buffer (140 mM KCl, 10 mM HEPES, pH 7.4). Injected cells were incubated at 37°C for the indicated times, then fixed with 4% paraformaldehyde in 1X PBS for 20 minutes at 4°C.

### Fluorescence in situ hybridization (FISH) and oligo-dT staining

For mRNA staining, cells were washed twice in 1X Sodium Saline Citrate (SSC; 150mM NaCl, 15mM sodium citrate, pH 7.0) buffer supplemented with 60% formamide. Cells were then treated with hybridization buffer (60% formamide, 100 mg/ml dextran sulfate, 1 mg/ml yeast tRNA, 5 mM vanadyl riboside complex, 1X SSC) containing 200 nM Alexa 546-conjugated ssDNA probe (Integrated DNA Technologies) for 18-24 hours at 37°C . Subsequently, coverslips were washed with 1X SSC supplemented with 60% formamide and the coverslips were mounted on DAPI.

The probe oligonucleotide sequences included:

anti-ftz (GTCGAGCCTGCCTTTGTCATCGTCGTCCTTGATAGTCACAACAGCCGGGACAACAC CCAT)

anti-βG (CTTCATCCACGTTCACCTTCGCCCCACAGGGCAGTAACGGCAGACTTCTCCTCAGGA GTCA).

For poly(A) mRNA staining, cells were washed in 1X SSC supplemented with 25% formamide. Cells were treated with poly(A) hybridization buffer (100 mg/ml dextran sulfate, 1 mg/ml yeast tRNA, 5 mM vanadyl riboside complex, 1X SSC) containing 200nM Alexa 546-conjugated ssDNA 60mer dT oligonucleotide (Integrated DNA Technologies).

### Stellaris single-molecule (sm)FISH

Stellaris smFISH experiments were performed as previously described (PMID: 32710402). Following fixation with 4% paraformaldehyde (Electron Microscopy Sciences), coverslips were submerged in methanol at −20°C for 30 minutes and then rehydrated twice in 1X PBS, each time incubating for 15 min. Cells were then washed twice in 1× SSC buffer (150 mM NaCl, 15 mM NaCitrate, pH 7.1) with 10% formamide, and incubated with 100 μl of hybridization buffer (100 mg/ml dextran sulfate, 10–15% formamide, 2× SSC) containing 125 nM Stellaris probes (LGC Biosearch Technologies) for 48 h.

### Immunofluorescence

Following fixation with 4% paraformaldehyde, the coverslips were washed with 1X PBS twice then incubated with 0.1% TritonX-100 in 1X PBS for 15 minutes. Subsequently, the coverslips were washed with 1X PBS twice, and incubated with either anti-TIA-1 antibodies (mouse monoclonal, 1:200 dilution, Invitrogen), anti-G3BP1 (mouse monoclonal, 1:500 dilution, Abcam), anti-FMRP (rabbit polyclonal, 1:50 dilution, Cell Signaling Technology), anti-PABP (rabbit polyclonal, 1:200 dilution, Abcam), anti-Caprin (rabbit polyclonal, 1:200 dilution, Proteintech) anti-Dcp1a (mouse monoclonal, 1:200 dilution, Sigma), and anti-P-eIF2α (rabbit polyclonal, 1:200 dilution, Cell Signaling Technology) antibodies for 1 hour. Antibody incubation was supplemented with 0.1mg/ml RNase-free BSA and 0.1% TritonX-100 in 1X PBS. In between antibody incubations, coverslips were washed by being placed cell-side down onto 0.5 mL droplets of 1X PBS on parafilm and allowed to incubate at room temperature three times for 5 minutes. Lastly, the coverslips were mounted onto glass coverslips using DAPI Fluoromount-G stain mounting solution (Southern Biotech).

### Imaging and image analysis

Cells were imaged using a fluorescence microscope (Nikon). Image analysis, including the stress granule assembly, eIF2α phosphorylation and Pearson’s correlation analysis, was performed using Nikon Imaging Software (NIS) Elements Advanced Research (Nikon). For stress granule assembly analysis, cells containing one or more stress granules were identified as stress-granule-positive cells. For eIF2α phosphorylation analysis, whole cell intensity for P-eIF2α staining was measured and normalized to cell-absent background intensity. For mRNA colocalization analysis, rectangular regions of interest were drawn to cover a single mRNA-positive structure in the cytosol and its surroundings (1‒4 μm^2^). The Pearson correlation coefficients between mRNA FISH and TIA-1/G3BP1/Dcp1a immunofluorescence was calculated by NIS analysis software. Examples are shown in Figure 1E. For each experiment, a total of 150-200 mRNA-positive structures across multiple cells in the cytosol were analyzed. The Pearson correlation between the mRNA microinjected in the cytosol and nuclear DAPI staining was also assessed to determine background levels of correlation.

### Immunoblotting

For immunoblot analysis, U2OS cells were washed twice with ice-cold 1X PBS, then lysed with sample buffer containing 50 mM Tris-HCl (pH 6.8), 2% SDS, 10% glycerol, 0.1% bromophenol blue and 1% β-mercaptoethanol, pre-heated at 95°C. The whole-cell lysates were subsequently further heated at 95°C for 5 minutes and then rested on ice for 5 minutes. The lysate was passed through –gauge needle to shear genomic DNA. Whole-cell lysates were separated by SDS-PAGE on 6–15% polyacrylamide gels, transferred to nitrocellulose membranes and probed with primary antibodies against PKR (rabbit monoclonal, 1:3000 dilution, Abcam), phospho-PKR (T446) (rabbit monoclonal, 1:3000 dilution, Abcam), phospho-eIF2α (rabbit polyclonal, 1:2000 dilution, Cell Signaling Technology), and tubulin (mouse monoclonal DM1a, 1:1000 dilution, Sigma)). Chemiluminescence luminol reagent (Pierce) and the Versadoc system (Bio-Rad) were used to visualize the blots.

### Puromycin, homoharringtonine, sodium arsenite treatments

U2OS cells were plated on 22 × 22 mm coverslips (VWR) in 35 mm mammalian tissue culture dishes (Thermo Scientific) for 24 h prior to treatment or transfection. Cells were subsequently treated with DMSO, puromycin (1 mM final), homoharringtonine (5 μM final), or sodium arsenite (100, 50, 25, 12 μM final).

### *In vitro* poly(I:C) transfection

Synthetic dsRNA analogue poly(I:C) HMW (Invivogen) was transfected into U2OS cells using JetPRIME transfection reagent (Roche Applied Science, Mannheim, Germany), according to the manufacturer’s instructions.

## Acknowledgments

We thank P. Ivanov and P. Andersen for providing us with the ΔΔG3BP1/2 MEFs and R. Mazroui for providing the eIF2α S51A MEFs. We thank members of the Lee and Palazzo labs and Carol Featherstone for their feedback in preparation with this manuscript. This work was supported by a grant to AFP from Natural Sciences and Engineering Research Council of Canada (FN 492860) and a grant to HOL from Canadian Institutes of Health Research (PJT-175174) and funding from Canada Research Chairs Program. The authors declare no competing financial interests.

## Author contributions

Conceptualization, S.J.I, A.F.P., H.O.L.; Funding and Supervision, A.F.P., H.O.L.; Investigation, S.J.I.; Analysis, S.J.I, L.F-W., A.F.P., H.O.L; Writing – Original Draft, S.J.I, A.F.P., H.O.L.; Writing – Review and Editing, S.J.I, A.F.P., H.O.L.

---

**Supplemental Figure 1.**
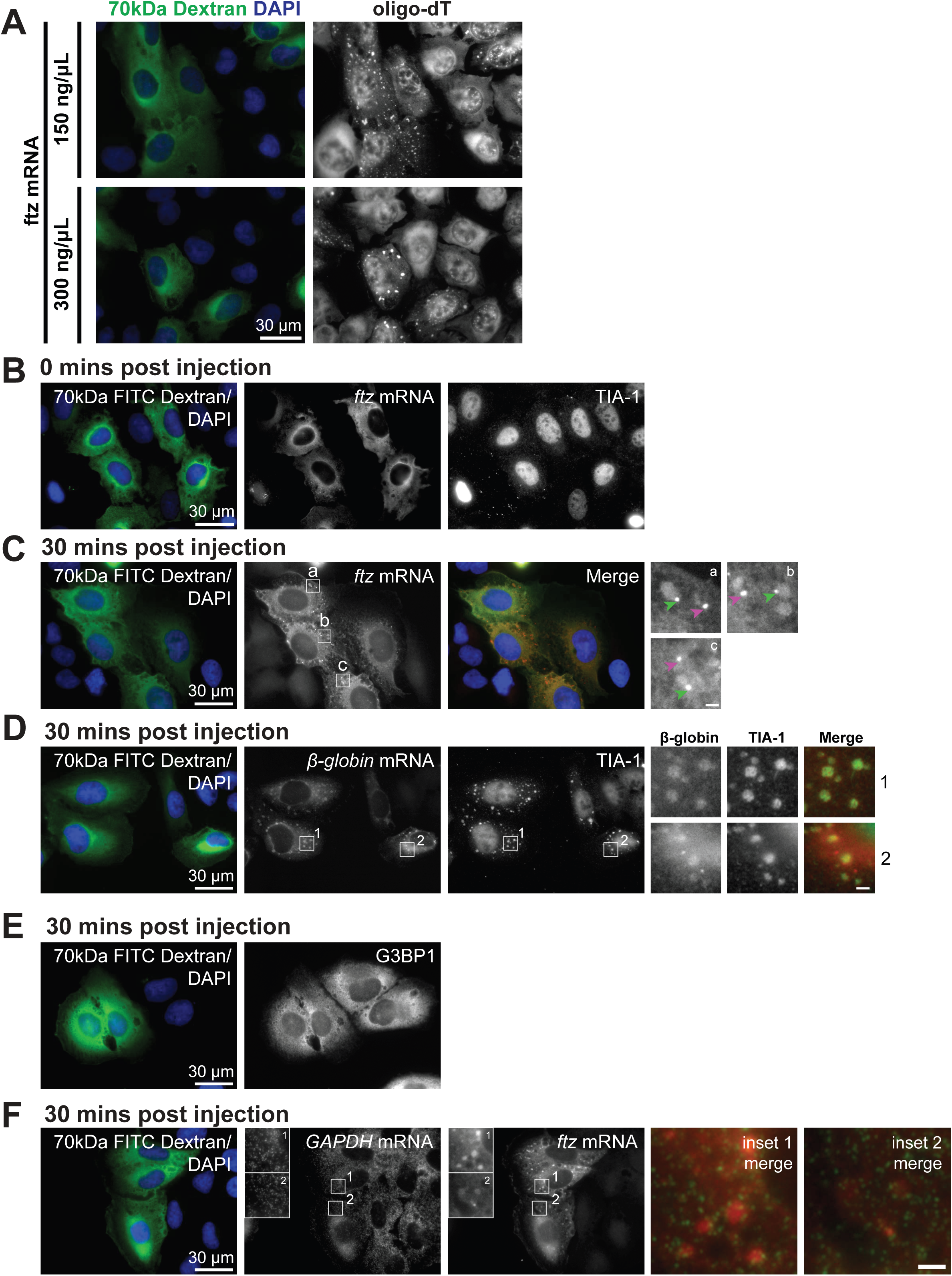
Related to Figure 1. **(A)** U2OS cells were microinjected with *ftz* mRNA (300 or 150ng/μL) and 70kD dextran conjugated to FITC to mark the microinjected cells and the compartment (cytoplasm). 30 minutes after microinjection, the cells were fixed and stained against poly(A)-RNA by using oligo-dT probes, and DNA by DAPI. All panels in each row represent a single field of view imaged for the indicated components. Scale bars: whole cell=30µm. **(B)** U2OS cells were microinjected with *ftz* mRNA (300ng/μL) and 70kD dextran conjugated to FITC to mark the microinjected cells and the compartment (cytoplasm). Immediately after microinjection (C) or after 5 minutes (D), the cells were fixed and stained against *ftz* mRNA using FISH probes, the SG-marker TIA-1 by immunofluorescence, and DNA by DAPI. Scale bars: whole cell=30µm. **(C)** U2OS cells were microinjected with *ftz* mRNA (300ng/μL) and 70kD dextran conjugated to FITC. After 30 minutes the cells were fixed and stained with FISH probes against *ftz* mRNAs and DAPI to stain DNA. All panels represent a single field of view imaged for the indicated components with an overlay, where FITC-70kD dextran is in green, *ftz* mRNA is in red and DAPI in blue. Zoomed in images are shown in boxed insets on the right. Note the large diffuse mRNA aggregates and the small mRNA foci. Scale bars: whole cell=30µm, insets=2µm. **(D)** U2OS cells were microinjected with *β-globin* mRNA (300ng/μL) and 70kD dextran conjugated to FITC. After 30 minutes, the cells were fixed and stained against *β-globin* mRNA using FISH probes, the SG-markers G3BP1 and TIA1 by immunofluorescence, and DNA by DAPI. All panels represent a single field of view imaged for the indicated components. Zoomed in images are displayed in boxed insets (1). The merged insets show *β-globin* mRNA in red and TIA-1 in green. Scale bars: whole cell=30µm, insets=2µm. **(E)** U2OS cells were microinjected with 70kD dextran conjugated to FITC alone. After 30 minutes the cells were fixed and stained against the SG-marker G3BP1 by immunofluorescence.Scale bars: whole cell=30µm. **(F)** U2OS cells were microinjected with *ftz* mRNA (300ng/μL) and 70kD dextran conjugated to FITC. After 30 minutes the cells were fixed and stained against *ftz* mRNA using FISH probes and endogenous *GAPDH* mRNA using single-molecule FISH probes, and DNA by DAPI. Scale bars: whole cell=30µm, insets=2µm. Zoomed in images are shown in boxed insets. The merged insets show *GAPDH* mRNA in green and *ftz* mRNA in red.

**Supplemental Figure 2.**
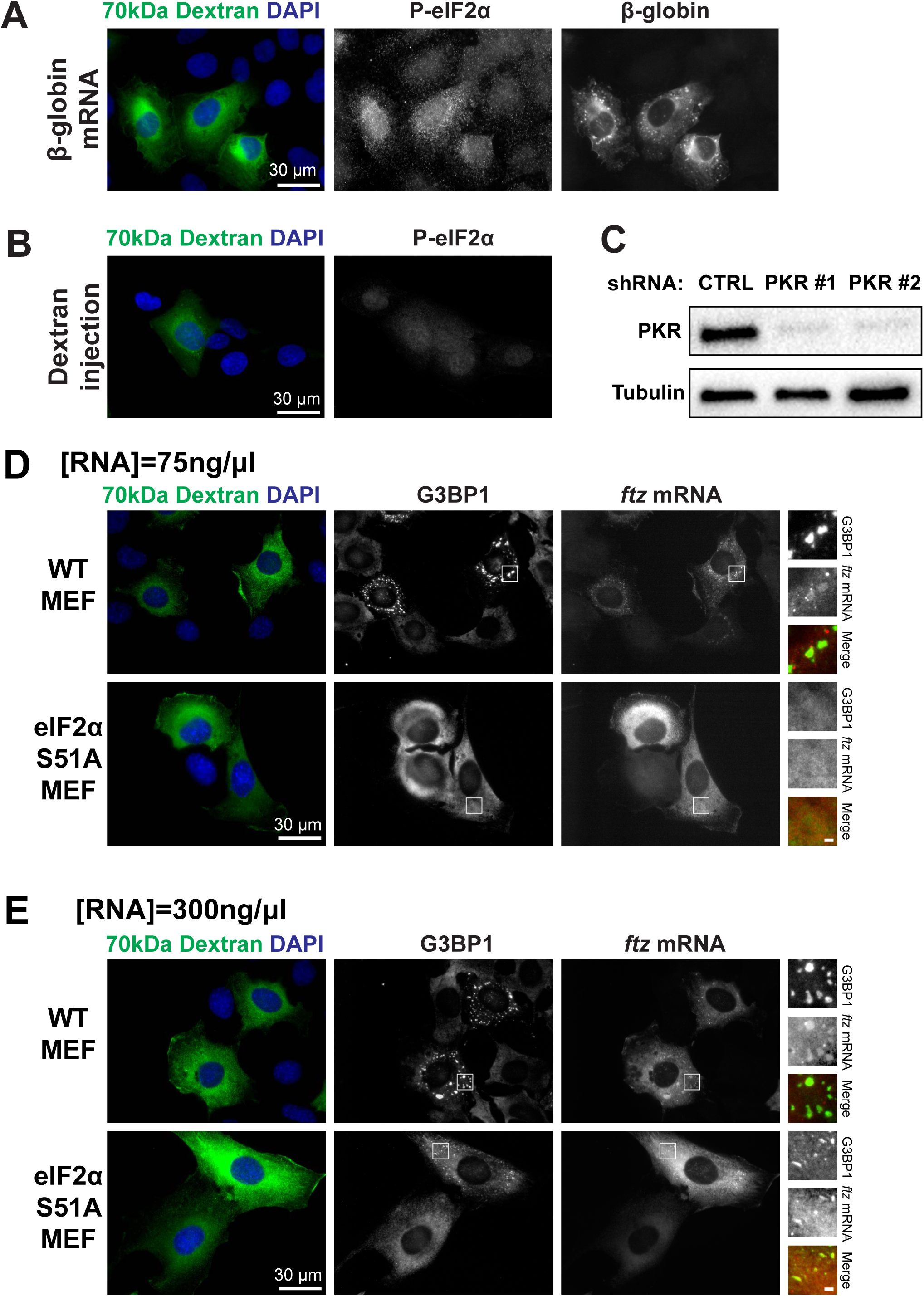
Related to Figure 2. **(A)** U2OS cells were microinjected with *β*-*globin* mRNA (300ng/μL) and 70kD dextran conjugated to FITC. After 30 minutes, the cells were fixed and stained against *β-globin* mRNA using FISH probes, and phospho-eIF2α by immunofluorescence, and DNA by DAPI. **(B)** U2OS cells were microinjected with buffer containing 70kD dextran conjugated to FITC. After 30 minutes, the cells were fixed and stained phospho-eIF2α by immunofluorescence and DNA by DAPI. **(C)** U2OS cells were transduced with lentivirus delivered shRNAs against PKR (two clones were used, PKR #1 and PKR #2) or a scrambled shRNA. After 2 days, the cell lysates were collected and separated by SDS-PAGE and immunoprobed for PKR and α-Tubulin, which served as a loading control. **(D)** Wild-type or eIF2α S51A phosphodeficient MEF cells were microinjected *ftz* mRNA (75ng/μL) and 70kD dextran conjugated to FITC, fixed 30 minutes after injection, and stained against *ftz* mRNA by FISH, G3BP1 by immunofluorescence, and DNA by DAPI. Scale bars: whole cell=30µm, insets=2µm. Zoomed in images are shown in boxed insets. The merged insets show G3BP1 in green and *ftz* mRNA in red. **(E)** Wild-type or eIF2α S51A phosphodeficient MEF cells were microinjected ftz mRNA (300ng/μL) and 70kD dextran conjugated to FITC, fixed 30 minutes after injection, and stained and stained against *ftz* mRNA by FISH, G3BP1 by immunofluorescence, and DNA by DAPI. Scale bars: whole cell=30µm, insets=2µm. Zoomed in images are shown in boxed insets. The merged insets show G3BP1 in green and *ftz* mRNA in red.

**Supplementary Figure 3.**
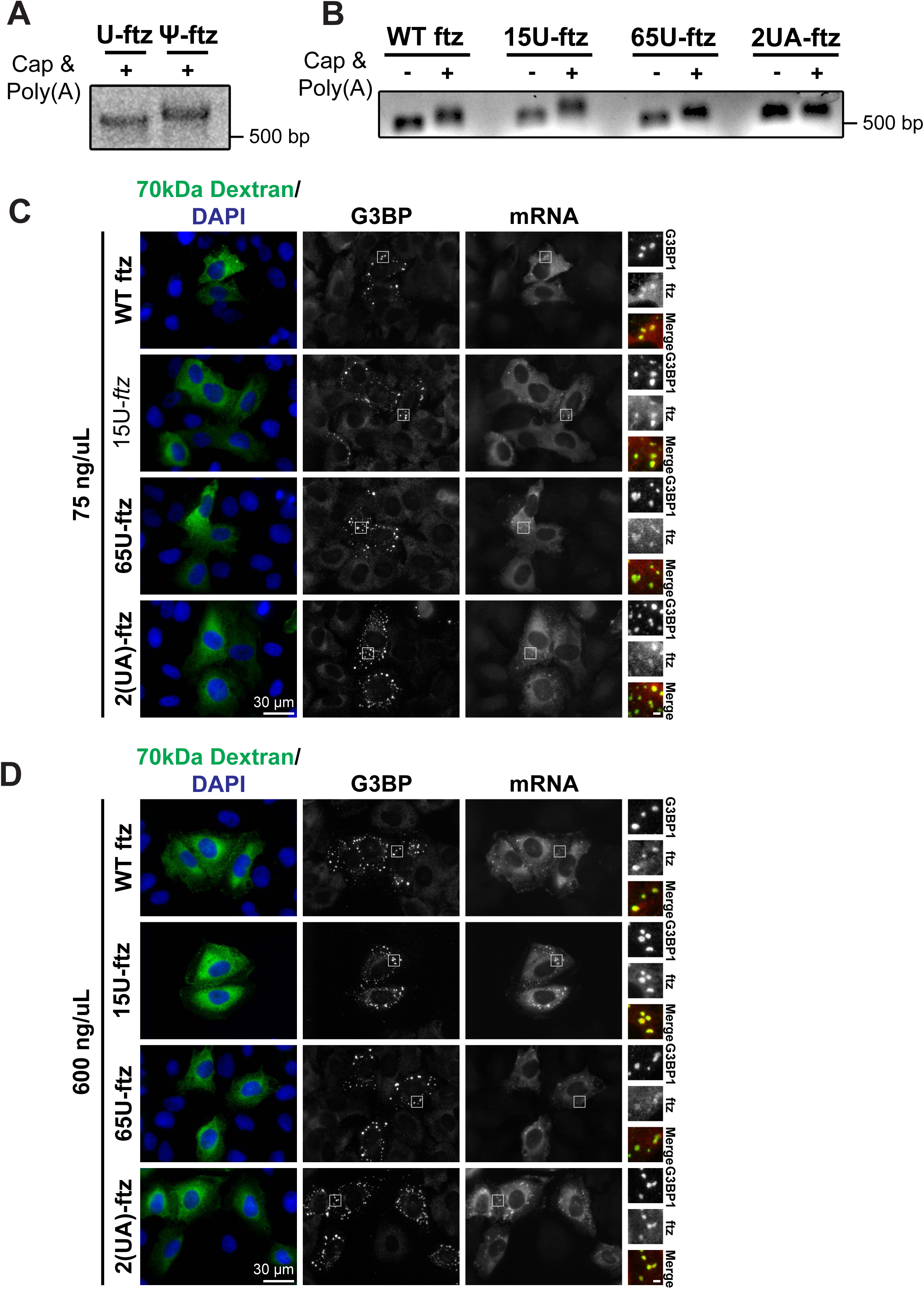
Related to Figure 3. **(A-B)** *ftz* mRNAs of varying nucleotide composition were *in vitro* transcribed, capped and polyadenylated, then purified. Samples of mRNAs before (“-”) and after (“+”) capping and polyadenylation were collected and resolved on an a 1% agarose gel. The migration of standards are indicated on the right. *ftz* mRNAs transcribed with pseudouridine as a replacement for uridine (“ψ-*ftz*”) are shown in (A), compared to wild-type uridine-containing *ftz* mRNA (“U-*ftz*” or “WT-*ftz*”). WT-, 15U-, 65U-, or 2(UA)-*ftz* mRNAs are shown in (B). **(C-D)** U2OS cells were microinjected with WT-, 15U-, 65U-, or 2(UA)-ftz mRNA at 75ng/μL (C) or 300ng/μL (D) and 70kD dextran conjugated to FITC. After 30 minutes, the cells were fixed and stained against ftz mRNA using FISH probes, the SG-markers G3BP1 by immunofluorescence, and DNA by DAPI. All panels in each row represent a single field of view imaged for the indicated components. Scale bars: whole cell=30µm, insets=2µm. Zoomed in images are shown in boxed insets. The merged insets show G3BP1 in green and *ftz* mRNA in red.

**Supplementary Figure 4.**
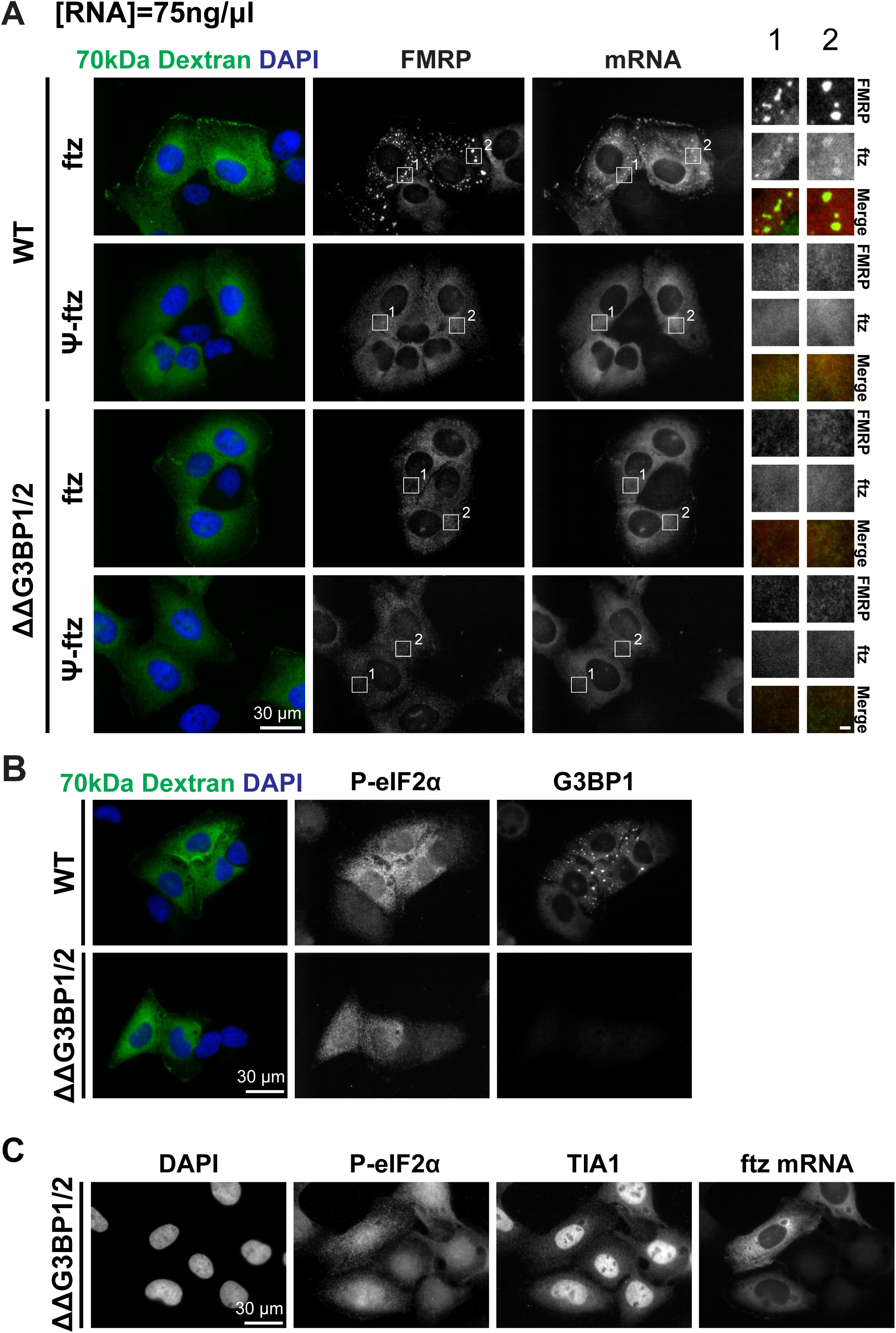
Related to Figure 5. **(A)** WT and ΔΔG3BP1/2 U2OS cells were microinjected with uridine-containing (“U-*ftz*”) or pseudouridine-containing *ftz* mRNA (“ψ-*<u>ftz</u>*”) at 75ng/μL and 70kD dextran conjugated to FITC. After 30 minutes, cells were fixed and stained for ftz mRNA using FISH probes, the SG-marker FMRP by immunofluorescence, and DNA by DAPI. Scale bars: whole cell=30µm, insets=2µm. Zoomed in images are shown in boxed insets. The merged insets show G3BP1 in green and *ftz* mRNA in red. **(B)** WT and ΔΔG3BP1/2 U2OS cells were microinjected with ftz mRNA at 300ng/μL and 70kD dextran conjugated to FITC. After 30 minutes, cells were fixed and stained for G3BP1 and phospho-eIF2α by immunofluorescence, and DNA by DAPI. Scale bars: 30µm. **(C)** ΔΔG3BP1/2 U2OS cells were microinjected with ftz mRNA as shown in (A), fixed 30 minutes after injection, and stained for ftz mRNA using FISH probes, TIA-1 and phospho-eIF2α by immunofluorescence, and DNA by DAPI. Scale bars: 30µm.

